# Behavioral encoding across timescales by region-specific dopamine dynamics

**DOI:** 10.1101/2022.12.04.519022

**Authors:** Søren H. Jørgensen, Aske L. Ejdrup, Matthew D. Lycas, Leonie P. Posselt, Kenneth L. Madsen, Lin Tian, Jakob K. Dreyer, Freja Herborg, Andreas T. Sørensen, Ulrik Gether

**Author notes:** These authors contributed equally to the work (Søren H. Jørgensen, Aske L. Ejdrup). **Author contributions:** S.H.J, A.L.E., A.T.S and U.G. conceptualized the study. L.T. and J.K.D. provided methodology. S.H.J. established the fiber photometry platform. S.H.J. performed surgeries and experiments with help from L.P.P. A.L.E. developed the analyses. A.L.E. and M.D.L. designed animal tracking and behavioral prediction. A.L.E. and S.H.J. analyzed the data. S.H.J, A.L.E, U.G., A.T.S, F.H.J, K.L.M., M.D.L and L.P.P. interpreted the data. A.L.E. drafted the manuscript. A.L.E., S.H.J. and U.G. edited the manuscript. All authors reviewed and critically evaluated the manuscript. **Competing interests Statement:** L.T. has ownership interests (stock, stock options, royalty, receipt of intellectual property rights/patent holder, excluding diversified mutual funds) and is a co-founder of Seven Biosciences. The other authors declare no competing interests.

## Abstract

The dorsal (DS) and ventral striatum (VS) receive dopaminergic projections that control motor functions and reward-related behavior. It remains poorly understood how dopamine release dynamics across different temporal scales in these regions are coupled to behavioral outcomes. Here, we employ the dopamine sensor dLight1.3b together with multi-region fiber photometry and machine learning-based analysis to decode dopamine dynamics across striatum during self-paced exploratory behavior in mice. Our data show a striking coordination of rapidly fluctuating signal in the DS, carrying information across dopamine levels, with a slower signal in the VS, consisting mainly of slow-paced transients. Importantly, these release dynamics correlated with discrete behavioral motifs, such as turns, running and grooming on a subsecond-to-minutes time scale. Disruption of dopamine dynamics with cocaine caused randomization of action selection sequencing and disturbance of DS-VS coordination. The data suggest that distinct dopamine dynamics of DS and VS jointly encode behavioral sequences during unconstrained activity with DS modulating the stringing together of actions and VS the signal to initiate and sustain the selected action.

**Significance Statement:** New genetically encoded dopamine sensors offer unprecedented temporal resolution for measurement of dopamine release dynamics across different brain regions over extended periods. In this study, we use the dopamine sensor dLight1.3b to decipher the role of dopamine release dynamics in the dorsal (DS) and ventral striatum (VS) of mice during simple, self-paced exploratory behavior. By AI-based splitting of behavioral kinematics into individual motifs, we link differential but highly cooperative dopamine release dynamics of DS and VS with movements on a subsecond-to-minutes time scales. In addition to coupling region-specific dopamine dynamics to behavioral sequences, our study demonstrates the strength of a machine learning-based data analysis pipeline that can be readily applied to other neurotransmitters for which genetically encoded biosensors are available.

## Introduction

It is essential to our survival that we can learn from experience and effectively select and sculpt our actions. By regulating reinforcement learning and movements, dopamine (DA) plays a fundamental role in this process (1–6). Interestingly, DA neurotransmission differs substantially from the classical fast synaptic transmission, as DA is a modulatory neurotransmitter exerting its effects largely via volume transmission (7, 8). DA is to a large extent released from non-synaptic varicosities to act on extrasynaptic metabotropic receptors situated on target neurons that can be micrometers away (9). Nevertheless, DA is believed to control a plethora of physiological functions with high precision across a broad range of temporal scales (9).

The dorsal (DS) and ventral striatum (VS) constitute major target areas for dopaminergic neurons. DS is heavily innervated by projections from substantia nigra pars compacta (SNc), while VS, including the nucleus accumbens, is innervated by the ventral tegmental area (VTA) (2, 5). Numerous studies have supported that DA DS projections predominantly play a role in locomotor control while DA VS projections mainly have been implicated in reward-related behavior (2, 4, 5, 10–15). It is well established that dysfunction of the DA system has major impact on striatal function and contributes to a range of disorders. Loss of nigrostriatal projections, together with impairment of motor functions, is a hallmark of Parkinson’s disease, while altered function of the VTA projections is thought to contribute to neuropsychiatric diseases like schizophrenia, depression and addiction (16–18).

Several methods have been used to study the firing and release patterns of DA neurons, such as fast-scan cyclic voltammetry (FSCV), microdialysis and electrophysiology (19–21). These techniques have revealed important functional characteristics of DA neurons, including “tonic” pacemaker-like firing and its “phasic” counterpart with trains of rapid depolarizations coordinated across neurons (3, 22–24). In response to unpredicted events, these phasic trains drive DA release on a sub-second to seconds time scale and is the putative driver behind reward-based learning. In contrast, it has been a widely held belief that motor function is modulated by slowly changing tonic activity on a time scale of tens of seconds to minutes (3). However, recent single unit recordings of nigrostriatal neurons have suggested that movement on a much faster time scale correlates with DA neuronal firing activity (25, 26). Similarly, measurements of Ca^2+^-activity in DA neurons support that sub-second changes in the activity of DA neurons may be important for triggering initiation of movement and possibly govern both future vigor and motor skill performance (11, 14, 27, 28). It is nevertheless largely unknown how these changes in neuronal activity are reflected in terminal DA release dynamics across various temporal and spatial scales in striatal subregions, and how these dynamics are linked to discrete behavioral outcomes during unconstrained activity in animals.

To address these questions, we set out to exploit recently developed genetically encoded biosensors for DA, representing an *in vivo* recording modality that enables real-time measurements of extracellular DA dynamics with millisecond temporal resolution over extended periods (29, 30). Using multi-region fiber photometry recording of dLight1.3b fluorescence, we investigate fast extracellular DA dynamics concurrently in DS and VS during self-paced exploratory activity. By employing extensive data analysis together with unsupervised machine learning algorithms, we stratify mouse behavioral kinematics into individual syllables that are further correlated across time scales with the highly distinct DA dynamics that we identify in the two regions. In summary, our data suggest that distinct DA dynamics of the DS and VS jointly orchestrate behavioral structure during spontaneous activity of the mice with a DS signal that modulates the stringing together of actions and a VS signal that provides the drive to initiate and sustain the action.

## Results

### Differential DA dynamics in DS and VS

To assess striatal DA dynamics in freely moving mice, we injected adeno-associated virus (AAV) encoding dLight1.3b (29) under control of the human synapsin promoter (AAV9-hSyn-dLight1.3b) in the right DS and left VS of tyrosine hydroxylase male (TH) Cre mice. The mice were also injected in the VTA with an AAV encoding Cre-dependent Gi-coupled DREADD (Designer Receptor Exclusively Activated by Designer Drugs, AAV8-hSyn-DIO-hM4Di-mCherry) to enable later chemogenetic inhibition of VTA DA neurons (SI Appendix, Fig. S1A). At least three weeks after surgery, we recorded dLight1.3b fluorescence via implanted optical fibers as a proxy for extracellular DA dynamics in the mice during self-paced unconstrained activity in an open-field arena (Fig. 1A and B, SI Appendix, Fig. S1B and S1C for calculation of zF). DS displayed incessant rapid fluctuations while VS showed slower and longer lasting fluctuations (Fig. 1B). These signals were interchangeable across hemispheres when sensor and fibers were shifted to left DS and right VS (SI Appendix, Fig. S1D). Note that these mice were not injected with AAV8-hSyn-DIO-hM4Di-mCherry into VTA, yet we still saw the same difference between DS and VS. Moreover, we observed no difference between the signal and isosbestic channel when mice were injected with GFP (AAV9-hSyn-EGFP-WPREpA) (SI Appendix, S1E) and found that the differences between DS and VS were patch cord fiber-independent (SI Appendix, S2A, B). Indeed, previous studies have also supported the strong specificity of the dLight sensors in DS and VS (see e.g. (29, 31, 32)). To analyze the recorded signals in greater detail, we computed the spectral energy density of the two regions, revealing a differential distribution of activity levels across frequency bands. Most of the DS signaling occurred in the 0.1-10 Hz range, whereas VS showed markedly slower dynamics (Fig. 1C, D). After photobleaching correction, DS showed minimal activity below 0.02 Hz. In contrast, VS exhibited DA fluctuations with durations upwards of 30 min but displayed minimal activity faster than 1.25 Hz (Fig. 1C, D and SI Appendix, Fig. S3A-C). Due to DS tapering off at 0.01 Hz, however, we decided to arbitrarily split the signal from both regions in two domains for later data interpretation purposes; a slow domain defined as fluctuations of 0.01 Hz and below, and a rapid domain defined as anything above (Fig. 1C and SI Appendix, Fig. S3A, B).

**Fig. 1.**
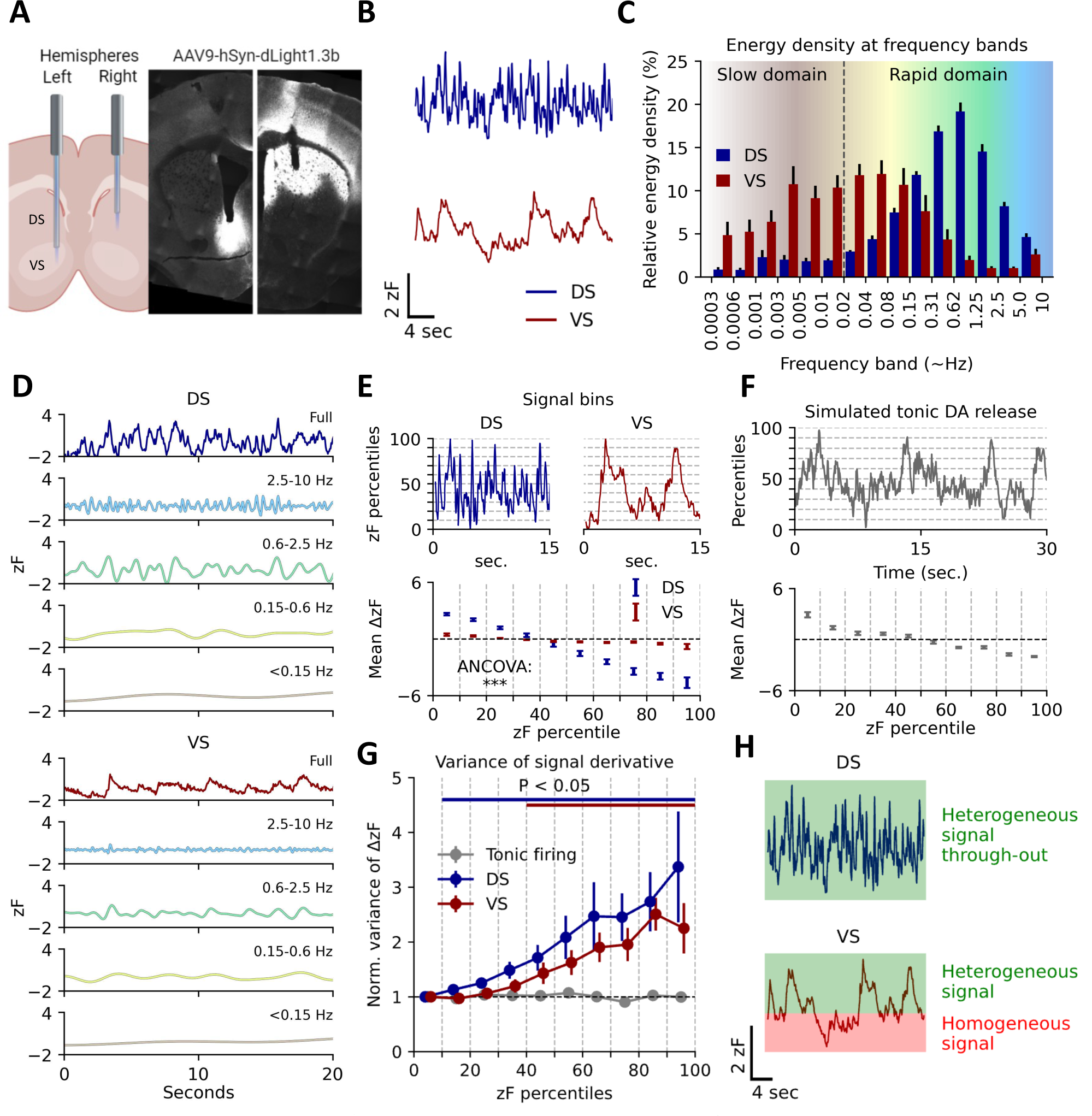
Basal DA dynamics differ between DS and VS during self-paced exploratory activity. (A) Injection of AAV9-hSyn-dLight1.3b in the right DS and left VS of Th-Cre mice followed by implantation of 200 µm optical fibers. Histology on coronal sections of the striatum show dLight1.3b expression and fiber location. (B) Representative traces from right hemisphere DS and left hemisphere VS DA fluctuations as assessed by dLight1.3b fluorescence measured during self-paced exploratory activity in an open-field arena that the mice had been exposed to 2-3 times before. (C) Spectral energy density of the two regions by frequency band. Error bars indicate S.E.M., n = 8 mice. (D) Representative frequency domains for DS and VS isolated by multiresolution analysis of the maximal overlap discrete wavelet transform. Colors match the corresponding domain in (C). (E) Top: Representative traces of VS and DS split into percentile bins of ten across fluorescence amplitudes. Bottom: First order derivative for each percentile bin of fluorescence amplitude for both regions. Error bars indicate S.E.M., n = 8 mice, ANCOVA, region:percentiles, F = 1199.5, p = 1.8E-16). (F) Top: Poisson-driven tonic DA simulation. Fluctuations are present despite the temporally uniform process. Dashed lines indicate amplitude percentiles. Bottom: First order derivative for each percentile bin of fluorescence amplitude. Error bars indicate S.E.M., n = 10 simulations. (G) Variance of the first order derivative for each percentile bin normalized to bottom 10^th^ percentile for simulated tonic release and DS and VS measurements. Error bars indicate S.E.M., p < 0.05 indicated by bars matching region color, student’s t-test, H_0_ = 1, multiple comparisons corrected for each percentile with Benjamini-Hochberg procedure (⍺ = 0.05). (H) Conceptual Figure of interpretation of data in (G).

To further investigate the biological significance of the observed DA dynamics, we computed the first order derivative (Δ) of each decile of the signals (Fig. 1E, top). Plotting the mean derivative (mean ΔzF) by signal amplitude (zF percentile) revealed a steeper slope for DS compared to VS (Fig. 1E, bottom). This is in line with a more rapidly fluctuating signal in the DS than in VS (Fig. 1D) and supports that DS is capable of both releasing and clearing DA more swiftly than VS as suggested previously (33). To test whether the data were in line with the tonic-to-phasic firing model of DA signaling (22, 24, 34), we simulated striatal DA dynamics based on tonic firing (2-10 Hz) of DA neurons. The extracellular DA level arising from release from a high number of tonically driven terminals can accurately be modeled by differential equations with a homogenous Poisson process as input (35–37). This mirrors a regular tonic activity with random phase shifts between neurons. Indeed, fluctuations are still present if DA release dynamics is simulated by such a process (Fig. 1F, top). However, fluctuations derived from a Poisson process produce a homogenous signal, as they represent random variations in an otherwise homogeneous input (35–37). Similarly, in the tonic-to-phasic firing model of DA signaling tonic firing is a homogenous signal, whereas the informational encoding is added by phasic firing, or lack thereof, which is a heterogeneous process. A feature of a system driven by a Poisson process is the uniform variance of the first order derivative across intensity levels (Fig. 1F, bottom). Therefore, we computed the variance of the derivative across signal intensity for DS and VS (Fig. 1G), which represents a measure of heterogeneity in the release patterns. Importantly, VS dynamics matched the tonic-to-phasic firing model of DA signaling, where biological information is encoded by phasic firing giving rise to transients (6). That is, we observed no significant change in variance for the lowest half of the DA signal as compared to the lowest ten percentiles, implying that the lower intensity-ranges represent random fluctuations from such a constant Poisson process. This suggests that a basal level of DA is sustained by pacemaker activity. Signal heterogeneity only seemed to be added in the upper ranges, where the variance of the derivative was larger, and the signal was dominated by transients conceivably representing novel information encoded by changes in firing rates. The same was not true for DS. Instead, variance increased from the lowest decile throughout (Fig. 1G), indicating fluctuations across all DA levels are driven by different upstream input phenomena, rather than constant release from tonic firing. The concept is schematized in Fig. 1H.

### Exploration and reward-seeking behavior promotes ramp-like DA release in VS but not DS

We investigated DA dynamics of DS and VS in two simple behavioral paradigms believed to implicate DA release. First, we assessed DA release correlates at entry to a novel environment by habituating mice to an open area for 15 minutes and then opening a gate to a small, adjacent novel chamber (Fig. 2A and SI Appendix Fig. S4A). The VS signal exhibited a fast, ramp-like increase in fluorescence, beginning immediately before entry to the novel chamber and returning to baseline after ∼20 sec. As expected, the mice also responded to new environment with an increase in locomotion that persisted after the increased VS DA signal had subsided (Fig. 2B). There was no significant change in DS at entry although the average signal across mice indicated a timing of transients (Fig. 2A and SI Appendix Fig. S4C). There was, however, a correlation between locomotion in the novel area and the DA response for both DS and VS supporting a role of the DA signal in coding of movement velocity (SI Appendix, Fig. S4D). The gate to the novel chamber was closed after 5 minutes, and the mice were left in the “old” arena for 30 minutes. We subsequently placed female-soiled bedding, known to be a source of intrinsically rewarding pheromones (38), in a second adjacent chamber and opened the gate (Fig. 2C). As the mice approached the bedding, the signal in VS ramped up as a conceivable response to the odor of the bedding (Fig. 2C, D and SI Appendix, Fig. S4E). When splitting the signal around the first encounter into frequency bands it was clear that this signal change in VS was mainly driven by frequencies slower than 0.6 Hz (Fig 2E). Although peaks of DA release in DS were observed surrounding the encounter, there were no significant changes in DS (Fig. 2C-E and SI Appendix, Fig. S4B and S4E). We next stimulated the inhibitory DREADD, hM4Di, that we had expressed in the VTA DA neurons of the dLight1.3b injected mice (SI Appendix, Fig. S1A). Indeed, we have previously demonstrated that stimulation of hM4Di in VTA DA neurons reduces exploratory locomotor activity and neuronal firing activity in *ex vivo* slice experiments (39). Administration of the DREADD agonist CNO (clozapine N-oxide) (2 mg/kg) caused a long-lasting decrease in the slow signal of VS with no significant decrease observed for DS (SI Appendix, Fig. S4F). Note that we observed a short-lasting increase in both DS and VS immediately after injection, likely because of handling as this was also observed for vehicle (SI Appendix, Fig. S4F). Detailed analysis revealed shifts also in the rapid dynamics of VS (SI Appendix, Fig. S4G) with reduced power in the 0.1 to 2 Hz range in response to CNO (Fig. 2I). Furthermore, CNO reduced the magnitude of the response in VS to female bedding, while in parallel the CNO-treated mice spent more time closing the distance to the bedding (Fig. 2F, G, H). Summarized, the data support the central importance of the VTA DA neurons for both slow and fast DA dynamics in VS and are consistent with the role of VS DA in reward-seeking behavior. We also expressed hM4Di in DA SNc neurons but detected no effect of CNO administration on the DA signal for neither DS nor VS despite histological confirmation of expression (SI Appendix, Fig. S4H). We have no immediate explanation for this observation but speculate that it relates to the previously reported higher D2 auto-receptor expression and stimulation in nigrostriatal SNc neurons as compared to VTA neurons, leading to saturated Gi-stimulation (40, 41).

**Fig. 2.**
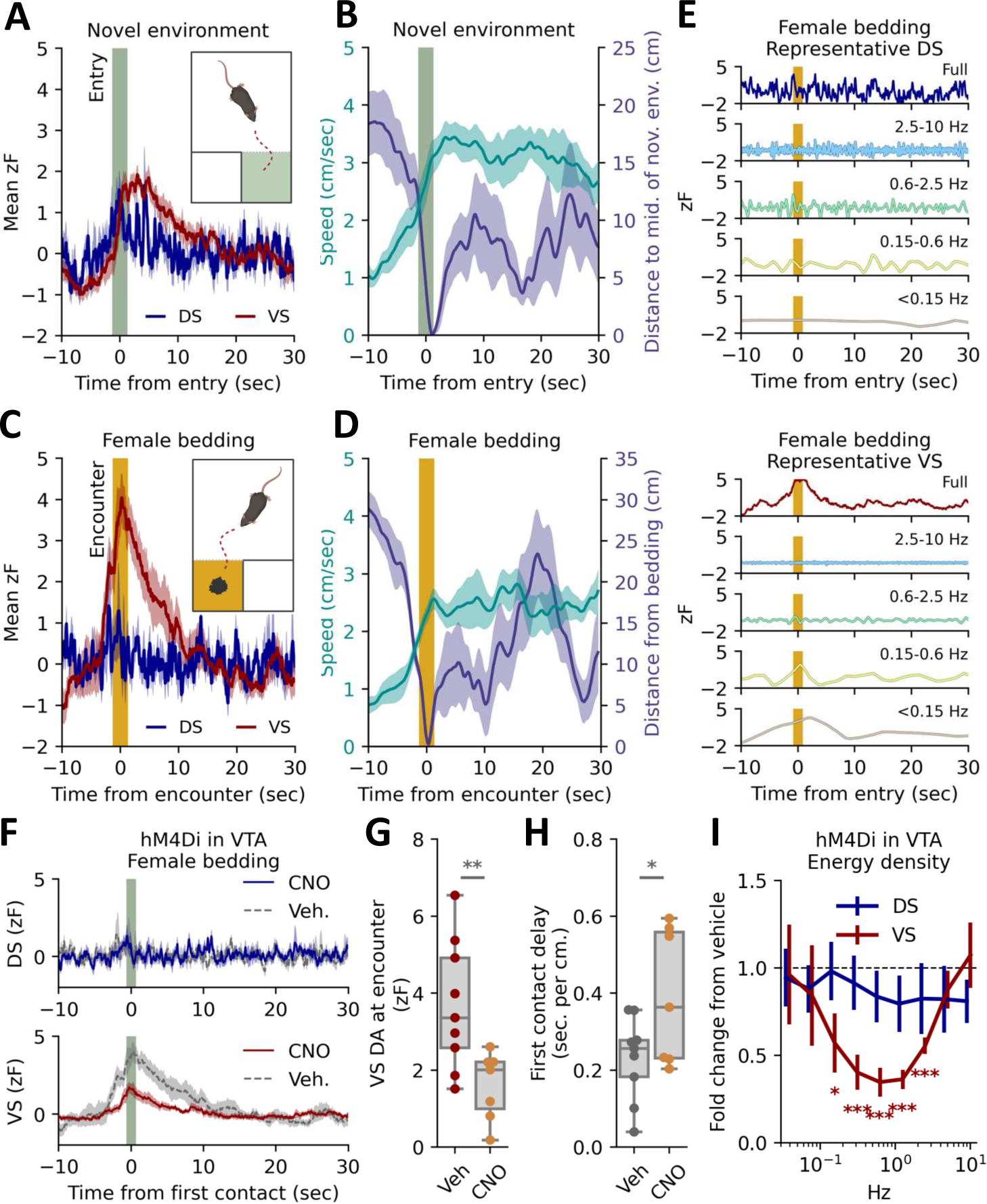
Exploration and reward-seeking behavior promotes ramp-like DA release in VS but not DS, which can be blunted by chemogenetic Gi-stimulation of VTA DA neurons. (A) Mean DA traces for right DS and left VS for TH-Cre mice expressing dLight1.3b in the right DS and left VS, and AAV-hSyn-DIO-hM4Di-mCherry in VTA. The traces are time-locked to entry into a novel chamber (t = 0, dark green band). Shaded area indicates S.E.M, n= 9 mice (two trials included for each mouse). (B) Mean speed (left y-axis) and distance to middle of novel chamber (right y-axis). Time-locked to entry (t = 0, dark green band). Shaded area indicates S.E.M, n= 9. (C) Mean fluorescent DA traces for right DS and left VS time-locked to first encounter with female-soiled bedding (t = 0, orange band). Shaded area indicates S.E.M, n= 9. (D) Mean speed (left y-axis) and distance to female bedding (right y-axis). Time-locked to first encounter (t = 0, orange band). Shaded area indicates S.E.M, n= 9. (E) DS (top) and VS (bottom) signals split into representative frequency domains isolated by multiresolution analysis of the maximal overlap discrete wavelet transform. Time-locked to first encounter with female-soiled bedding (t = 0, orange band). Colors match the corresponding domain in Fig. 1C. (F) Response to female bedding as in (c). after i.p. injection of either vehicle or 2 mg/kg CNO; DS (dark blue, top) and VS (dark red, bottom) signals are plotted for both vehicle (dashed line) and CNO (solid line). Shaded area indicates S.E.M., n = 9 for both groups). (G) Box plot of DA levels in VS at first encounter (**p = 0.009, unpaired student’s t-test, n = 9). (H) Box plot of delay from gate removed to first encounter – unit is sec/cm to account for differences in starting position for individual mice (*p = 0.04, unpaired student’s t-test, n = 9). (I) Change in spectral energy density from vehicle to CNO for minutes 15 to 30. Error bars indicate S.E.M., n = 9, student’s t-test, H_0_ = 1, FWER correction for all nine frequency bands with Bonferroni-Holm method, *p<0.05, ***, p<0.001.

### DA release in DS and VS correlates with action selection on a sub-second and minute-to-minute scale

The activity of D1R and D2R-expressing medium spiny neurons (MSNs) of the direct and indirect pathway, respectively, were recently shown to encode behavior within the DS by organizing moment-to-moment action selection with sub-second precision (42). An unresolved question is how this may be modulated by input from DA projections. We thus sought to determine how individual behavioral motifs, known as syllables (i.e., run, walk, turn etc.), might be encoded by DA release in DS and VS. Top-down video tracking of the behavior was analyzed using DeepLabCut (43) (Fig. 3A). Limb trajectories were tracked, segmented into individual syllables using the unsupervised clustering algorithm B-SOiD (44) and aligned with simultaneously recorded dLight1.3b DA traces (Fig. 3A, B). The clustering algorithm identified >50 unique syllables; however, a low usage and high inter-mouse variability was seen for the less frequent (SI Appendix, Fig. S5A), and consequently we focused on the twenty most recurring syllables (Fig. 3C). We computed the average DA trace at onset for each syllable from all identified instances in both regions (Fig. 3D). For several syllables we observed a distinct pattern in the averaged DA signal surrounding onset that differed between the two striatal regions. For DS, the general picture observed was fluctuations that peaked positively or negatively right at syllable onset. VS activity was also primarily fixed around syllable onset but changes were more gradual and longer lasting (Fig. 3D and SI Appendix, Fig. S5B and C).

**Fig. 3.**
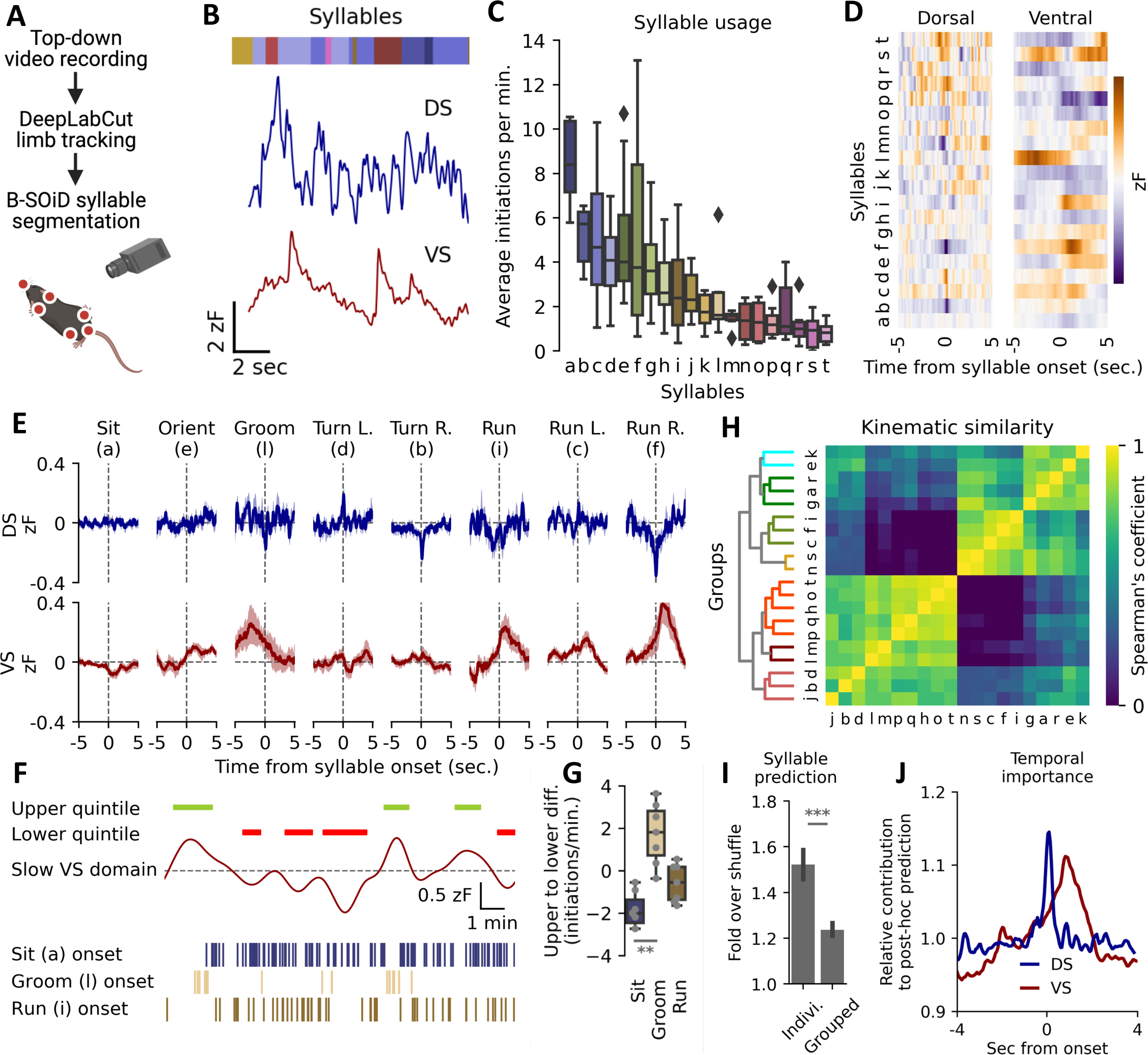
DS and VS DA dynamics correlate with movement across timescales. (A) Syllable identification pipeline. Mice were filmed from above, limbs tracked with DeepLabCut (43) and syllables segmented with an unsupervised clustering algorithm, B-SOiD (44). (B) Identified syllables (coloured bar, each colour representing a syllable) aligned to the DA signals in both regions. (C) Box plots of frequency of initiation for 20 most frequent syllables, n = 9. Syllables assigned letters in descending order. Black diamonds indicate outliers. (D) Average DA trace around onset in both DS and VS for each of the 20 most frequent syllables. (E) Fluorescent DA trace of DS (blue) and VS (red) around onset for eight humanly unambiguous syllables. Shaded area indicates S.E.M., n = 9. (F) Representative example showing frequency of sit (blue), groom (sand) and run (brown) correlated with slow DA oscillations in VS. Green bars indicate periods of upper quintile and red bars lower quintile. Syllable onsets are depicted with lines below. (G) Box plot of difference in frequency for upper and lower quintile of slow VS levels (one-way ANOVA for all 8 syllables in (E), F = 3.5, **p = 0.004. Post-hoc Tukey HSD: sit = groom, **p = 0.01, n = 9, see SI Appendix, Fig. S6A for statistics all 8 syllables). (H) Syllables hierarchically clustered by prediction certainty with Spearman correlation coefficient and segmented into seven differently coloured groups by arbitrary threshold (see SI Appendix, Fig. S5C for human annotations). (I) A random forest classifier correctly predicts individual syllables at a higher rate than kinematically grouped syllables when compared to shuffled guessing (right, fold over shuffle ± S.E.M., ***p = 5.2E-12, paired, two-tailed student’s t-test, n=9, 20 train-test iterations). (J) Relative contribution to classifier prediction by region (DS, blue. VS, red) around onset of syllables (n=9, 20 train-test iterations).

Eight syllables could unambiguously be classified by human observers, and the patterns of DA signal around onset are shown in Fig. 3E (see SI Appendix, Fig. S5B for all syllables). For two stationary syllables, sit and groom, we observed no detectable correlation to DS fluctuations. However, for grooming a distinct pattern emerged in VS with a transient increase in the signal peaking immediately before syllable onset, suggesting that the action either requires or promotes DA release. For turns, we observed mirrored DA activity for opposite turn directions. Ipsiversive (right) turns correlated with a brief depression at onset and contraversive (left) turns with a brief elevation in DA signal in right DS (Fig. 3E). Note that by expressing dLight1.3b and implanting fibers in the opposite hemispheres, we observed an identical mirrored pattern (SI Appendix, Fig. S6A). For run initiation, the most prominent feature was elevated DA levels in VS with no clear pattern for DS (Fig. 3E). Curiously, during running turns, we observed a pattern akin to the rise during a run in VS, accompanied by the DS pattern of a stationary turn, suggesting a modular design of at least some of the identified syllables (Fig. 3E). Note that as B-SOiD segments syllables at 100 ms resolution, peak activity timing during turns could not be temporally distinguished from onset (SI Appendix, S6B).

While the DS signal showed minimal minute-to-minute fluctuations, such slow fluctuations were present in VS (Fig. 1C and SI Appendix, Fig. S3B, C). Intriguingly, these slow fluctuations correlated with the propensity of using distinct syllables. Sitting was significantly more frequent during the bottom quintile of these slow fluctuations as compared to the top quintile, while grooming demonstrated the opposite pattern (Fig. 3F, G and SI Appendix, Fig. S6C). The run syllable, however, did not occur more frequently during long-term elevations, indicating a complex relationship between slow DA levels, second-to-second fluctuations and behavioral syllables.

We subsequently tested a two-part hypothesis; the DA traces predict movements, and motions with similar kinematic correlate with similar traces. We used Spearman’s correlation coefficient to cluster syllables by kinematics (Fig. 3H, see SI Appendix, Fig. S5D for human annotations) and we used machine learning by training a random forest classifier to predict syllables based on DA traces of both DS and VS. When predicting individual syllables, the classifier performed clearly better than chance when tested on a held-out data set (1.5-fold over shuffle, Fig. 3I) (see prediction for the eight most unambiguous syllables in SI Appendix, Fig. S5E), primarily using information at onset for DS and the seconds following for VS (Fig. 3J). In contrast, syllable prediction became less accurate (1.2-fold over shuffle) when predicting syllables grouped by kinematics (Fig. 3I). This highlights the predictive power of DS and VS DA traces when inferring behavior while also showing that movements of similar kinematics might differ substantially in their underlying DA signals.

### Cocaine alters rapid dynamics and disrupts DA correlation to action selection

Next, we investigated the effect of cocaine that inhibits DA reuptake by blockade of the dopamine transporter (DAT) (45). Cocaine (20 mg/kg i.p.) caused a substantial increase in the slow DA signal in both DS and VS, which as expected was accompanied by increased locomotor activity (Fig. 4A). No increase in fluorescent signal was observed in mice injected with AAV9-hSyn-EGFP-WPREpA (SI Appendix, Fig. S7A). The cocaine-induced increase in fluorescence in VS was 20-40 times the standard deviation of the signal at baseline, whereas slow DS fluctuations only increased with 4-8 standard deviations (SI Appendix, Fig. S7B). As the fluorescence is measured in arbitrary units and z-scored to baseline signal in each region, amplitudes in the two regions cannot be compared directly. Therefore, these observations may suggest that cocaine increases DA in VS to a greater extent as compared to DS or, alternatively, that the absolute DA fluctuations in DS under basal conditions are of a larger magnitude as compared to VS.

**Fig. 4.**
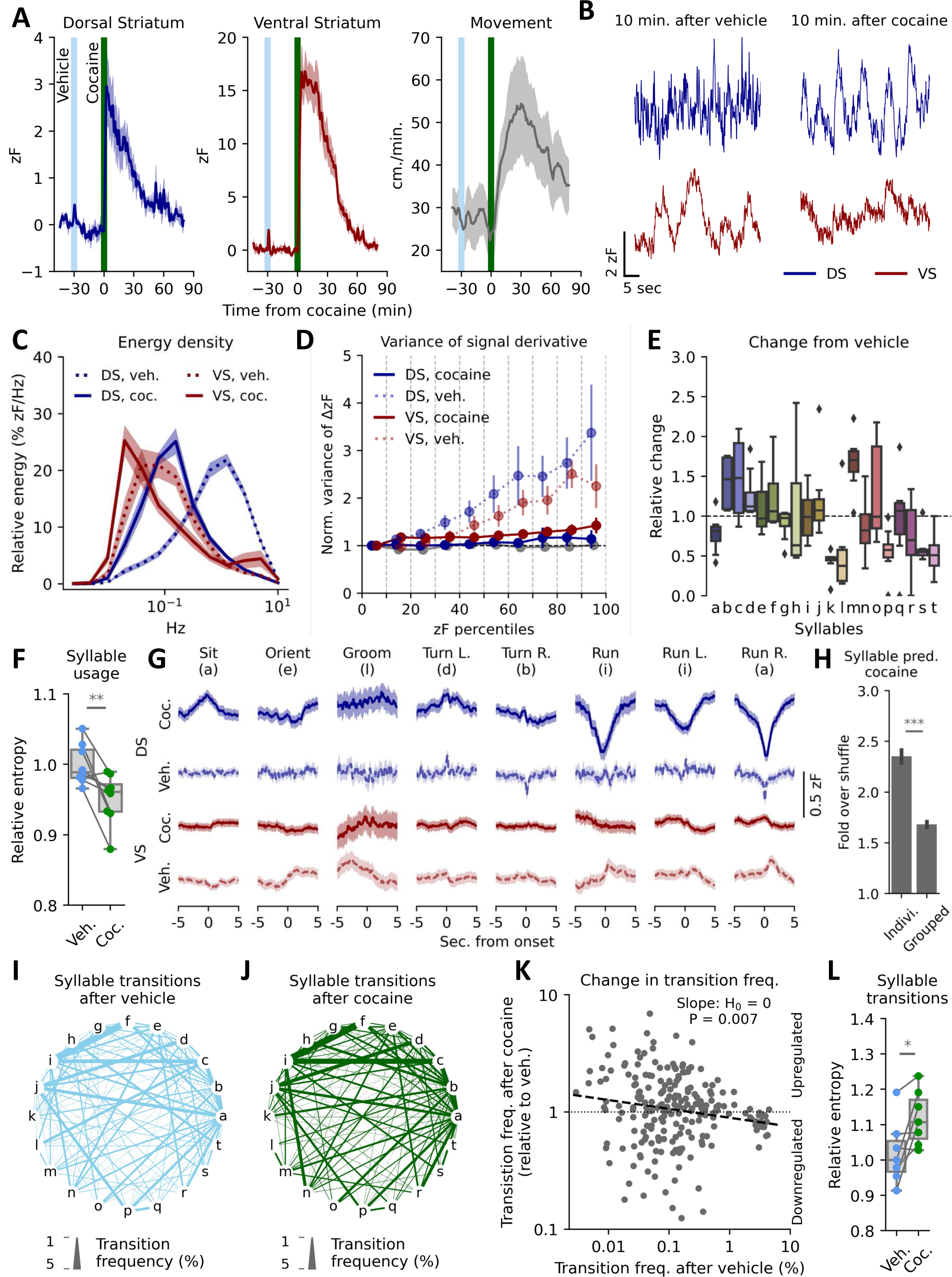
Cocaine alters rapid DA dynamics and ablates VS correlation to sub-second movements. (A) dLight1.3b expressing mice were habituated to an open arena for 15 minutes before receiving vehicle i.p., followed by 20 mg/kg cocaine after 30 min. Signal in DS (left) and VS (middle) and movement (right) is plotted in one-minute bins. Shaded areas indicate S.E.M., n = 8 mice. (B) Representative traces of rapid dynamics in DS and VS 10 min. after vehicle (left) and cocaine (right). (C) Spectral energy density of DS and VS signal 10 to 20 minutes after vehicle (dashes lines) and cocaine (solid lines). Shaded areas indicate S.E.M., n = 8 mice. (D) Variance of the first order derivative for each percentile bin normalized to bottom 10th percentile for simulated tonic release and DS and VS measurements. Error bars indicate S.E.M., no percentile survived multiple comparison with Benjamini-Hochberg procedure (⍺ = 0.05) after cocaine (student’s t-test, H0 = 1, n = 8). (E) Box plot of change in syllable frequency from vehicle to cocaine administration for the 20 most frequent syllables, n = 8 mice. (F) Entropy of syllable usage after vehicle (light blue) and cocaine (green). Normalized to vehicle mean (**p = 0.009, two-sided, paired student’s t-test, n = 8 mice). (G) Mean DA traces for DS (blue) and VS (red) after vehicle (dashed line) or cocaine (solid line) for eight selected syllables. Lines are averaged across mice with shaded areas indicating S.E.M., n = 8 mice. (H) A random forest classifier correctly predicts syllables 2-2.5 times as often as shuffled guessing and performs significantly better at individual syllables compared to kinematically similar groups (***p = 1E-21, paired, two-tailed student’s t-test). (I-J) State-transition plot of syllable transition frequency after vehicle (I) and cocaine (J) injection. Letters represent individual syllables with line thickness between the letters indicating transition frequency between the two syllables in absolute percentages. (K) Syllable transition frequencies after vehicle injection vs. transition frequency after cocaine relative to vehicle. Dashed line indicates least square linear fit (**p = 0.007, H_0_: slope = 0). (L) Entropy for the transition matrix (SI Appendix, Fig. S7E) after vehicle (light blue) and cocaine (green). Normalized to vehicle mean (*p = 0.015, one-sided, paired student’s t-test, n = 8 mice).

The fast DA dynamics changed radically 10 to 20 min after cocaine with a shift towards lower-frequency fluctuations (Fig. 4B, C and SI Appendix, Fig. S7C). The rapid VS signal was shifted so far towards low-frequency fluctuations that it started to blend with our threshold for slow fluctuations. This shift in VS was accompanied by a slight but significant increase in energy of the fastest domain (Fig. 4C and SI Appendix, Fig. S7C). Computing the variance of the 1^st^ derivative after vehicle resembled that found in DS and VS for untreated mice (Fig. 4D and Fig. 1G). In contrast, variance of the derivative changed drastically after cocaine to the point where no significant differences were observable across fluorescence intensities in DS and VS. This could suggest that the observed fluctuations after cocaine were random variations in a constant upstream process generated by a uniform input, such as tonic firing.

Analysis (as in Fig. 3) of behavioral syllables after cocaine showed a marked change in frequency of various syllables (Fig. 4E and SI Appendix, Fig. S7D, E). Syllable usage, however, did not become more disordered. Rather, entropy of the system decreased slightly (Fig. 4F), likely because of more frequent locomotion initiations. We also observed changes to the average DS for several syllables with run and run left/right being characterized by a strong depression of signal in DS around initiation. In contrast, sitting correlated with an uptick in DS release, as opposed to no correlation after vehicle injection. VS, on the other hand, lost most of its correlative signatures, which might be the result of the strongly elevated slow DA levels. Interestingly, the behavioral classifier became stronger at predicting syllables from DA traces after cocaine (Fig. 4H). This increase in predictive power came almost exclusively from the DS trace with a broader temporal window now contributing to the predictions (SI Appendix, Fig. 7F).

Because variance of derivative indicated that the observed DA fluctuations after cocaine were the result of random fluctuations in a homogenous signal, and because syllable usage did not appear more disordered, the DA fluctuations of DS did not seem to encode individual syllable selection. Rather, we hypothesized that the observed correlation between DA and syllables might reflect the stringing together of actions (4). To assess this, we mapped the frequency of transition from one syllable to another (Fig. 4I, J and SI Appendix, Fig. S7G). Interestingly, we observed that the most used transition frequencies after vehicle were downregulated after cocaine, whereas the less frequent transitions were generally upregulated (Fig. 4K). This corresponds to a decrease in transition orderliness as apparent when calculating the system entropy from the transition matrix (Fig. 4L and SI Appendix, Fig. S7E).

### DS and VS DA dynamics correlate on a sub-second and tens-of-second scale

Finally, we wanted to determine whether DA dynamics in DS and VS overall were temporally correlated. We therefore quantified the cross-correlation of the signals by time-shifting the DS trace (Fig. 5A). A maximal overlap for DA signal in the two regions was observed when DS was shifted ∼0.5 sec forwards (Fig. 5B), indicating that DA release in the DS is predictive of a subsequent VS release. DS continued to positively correlate with increased VS signal for up to 20 seconds. When shifting the DS signal backwards in time, we observed a negative correlation between DS and VS release from -10 to -20 seconds, showing that VS release predicted a future depression in DS signaling. This correlation was spread across frequency domains, with correlations across timescales (Fig. 5C).

**Fig. 5.**
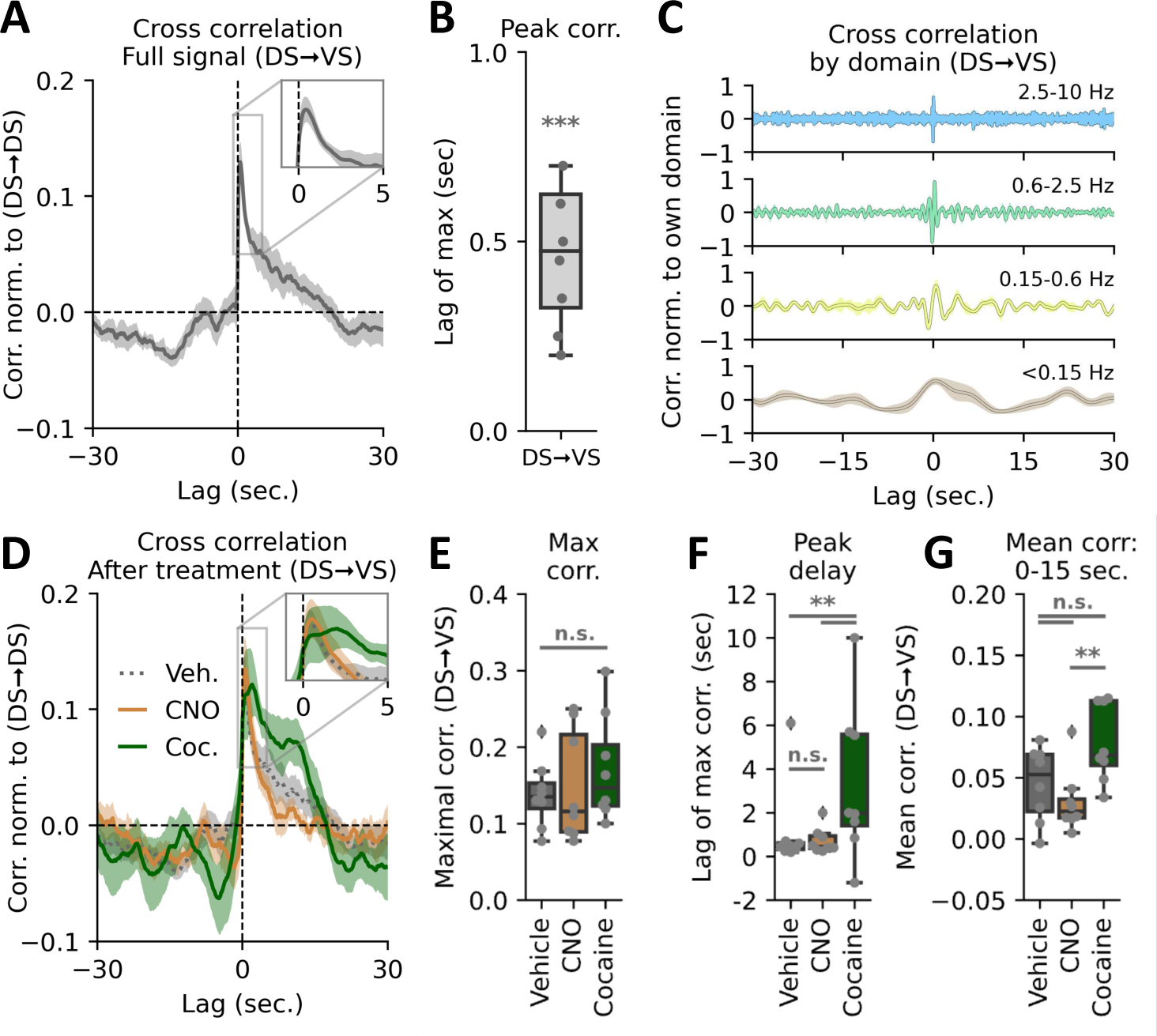
Both VTA inhibition and cocaine alters cross-correlation of intra-striatal DA activity. (A) DS-VS cross-correlation normalized to DS auto-correlation. Inset shows correlation peak and highlights that release in the DS is predictive of a subsequent VS release with approx. 0.5 sec lag time. Shaded area indicates S.E.M., n = 8 mice. (B) Box plot of lag of maximal DS-VS cross-correlation (***p = 4.4E-5, one-sample student’s t-test, n = 8 mice). (C) DS-VS cross-correlation by different frequency bands. Each domain is normalized to self within each mouse. Colours correspond to frequency domains in Fig. 1C. (D) DS-VS cross-correlation for vehicle (dashed grey line), CNO (solid orange) and cocaine (solid green). Shaded area indicates S.E.M., n = 8 mice for all three groups. (E) Maximal cross-correlation remains unchanged (one-way ANOVA: F = 0.58, p = 0.57, n = 8). All treatments normalized to DS auto-correlation. (F) Box plot of distribution of max lag in DS-VS cross-correlation for vehicle, CNO and cocaine. Only cocaine significantly alters correlation timing (veh.:CNO, p = 0.16; veh.:cocaine, **p = 0.002; CNO:cocaine, **p = 0.004, Levene variance test, FWER correction with Bonferroni-Holm, n = 8 mice). (G) Quantification of mean cross-correlation from 0 to 15 sec. VS lag. Vehicle is not significantly different from the two treatments, but CNO and cocaine differs (one-way ANOVA: F = 7.56, p = 0.0059, n= 8; Tukey HSD: veh.:CNO, p = 0.58; veh.:cocaine, p = 0.09; CNO:cocaine, **p = 0.01).

We subsequently assessed the correlation between the DS and VS signal in CNO-injected mice expressing hM4Di in VTA DA neurons. Strikingly, despite the major changes in VS dynamics (Fig. 2J), the peak correlation when normalized to DS autocorrelation was statistically identical to that of vehicle (Fig. 5D, E). Thus, metabotropic inhibition of VTA neurons is not sufficient to disturb the apparent coordination between the DS and VS DA signals. A different picture, however, emerged in mice upon cocaine exposure. As with CNO, the peak correlation when normalized to DS autocorrelation was not statistically different from vehicle (Fig. 5D, E). However, for cocaine we observed an altered lag of maximal cross-correlation after cocaine as compared to vehicle or CNO conditions with inter-mouse variation increasing significantly for cocaine (Fig. 5D, F). This suggests a temporal decoupling in the cross-regional coordination of DS and VS DA release. Moreover, the DS signal correlated with the VS signal further into the future for cocaine as compared to the VTA-inhibition by CNO (Fig. 5G). The difference between vehicle and the two treatments, however, was not significant.

## Discussion

The striatum constitutes an essential subcortical forebrain nucleus that receives major input from midbrain dopaminergic neurons and plays a key role in motor functions and reward-related behavior (2). Here, we use the recently developed, genetically encoded DA biosensor dLight1.3b to gain insights into how extracellular DA dynamics on different time scales in striatal subregions are linked to self-paced exploratory behavioral sequences. We achieve these insights by applying several machine learning-based data algorithms that enable highly detailed dissection of fast transmitter fluctuations in relation to discrete behavioral output. Together, our data show how major differences in the extracellular DA dynamics of the mouse DS and VS might underlie distinct, yet cooperative functions, of the two regions. Indeed, previous studies have indicated region-specific activity of DA neurons when the signals were aligned with well-defined behavioral events (11, 46–48). However, there has been little focus on subregion specific DA release dynamics over extended periods as we measure in the present study. Interestingly, the observed difference in sub-region-specific DA dynamics is not readily predicted from electrophysiological recordings of dopaminergic neuronal firing as these have reported similar overall firing rates of DA neurons in SNc and VTA (49–51). Thus, the differential DA release in substriatal regions may not be simply explained by differences in the neuronal firing rate, consistent with the recently demonstrated partial dissociation of DA release in VS from neuronal firing in the VTA (13).

Earlier studies involving primarily electrophysiological recordings and FSCV led to the classical assumption that bursts of DA release on a sub-second to second time scale is critical for reward-based learning while slower changing tonic activity is critical for control of movements (3). However, recent employment of electrophysiology and genetically encoded Ca^2+^ sensors to assess firing activity of DA neurons have implicated fast changes in activity in movement control (11, 14, 25–28). In the present study, we take this a step further to correlate highly differential DA fluctuations in DS and VS with discrete behavioral actions. By splitting the activity of the mice into behavioral syllables we show that initiation of individual movements, such as running and turning, correlate with increases or depressions in DA of sub-second to seconds duration in both DS and VS. Interestingly, the fast DA fluctuations observed in the DS may couple directly to the Ca^2+^-activity patterns of D1R- and D2R-expressing MSNs. In parallel to our findings, an increase in Ca^2+^-activity was observed for D1R- and D2R-expressing MSNs at onset of a contraversive turn while a decrease was seen for an ipsilateral turn (52). Indeed, this mirroring of the DA signal in Ca^2+^ dynamics of DA target neurons indicates that DA signaling in the DS on a hundreds-of-millisecond scale is critical for coordination of moment-to-moment movements controlled by the basal ganglia. Recently, ipsiversive depressions and contraversive increases were also observed at onset of turns in the DS by use of the genetically encoded sensor GRAB_DA_ (53), however, the changes observed lasted several seconds after onset which may be attributed to the slower off-rate kinetics of this sensor (53, 54).

While the DS signal correlated with movement on a fast timescale, syllable initiation rather coincided for the VS signal with second-long changes and slow fluctuations. Specifically, we observed changes in DS signal right at movement onset, whereas VS signal mainly increased immediately after initiation of high-velocity movements. This was observed in combination with the lateral pattern in DS during running turns, suggesting an element of modularity in the system. Moreover, our cross-correlation analysis supported considerable intra-striatal predictive power of the DA signal; hence, DA appears to be released in a coordinated and differential fashion to integrate the distinct functions of striatal subregions on sub-second to seconds time scales. Importantly, the data lend support to suggested interdependency proposed by Kelly & Moore in 1976 between VS and DS for executing motor behaviors such as turning (55). It may also be considered to what degree our findings may relate to recent observations indicating wave-like DA activation patterns across striatal regions (56). However, we cannot, at this stage, say how the cooperativity between DS and VS is coordinated or whether it is direct or indirect. We should note that we cannot, based on the current data, say whether the correlation between DS and VS only exist cross-hemispherically, as space constraints did not allow simultaneous measurements in DS and VS of the same hemisphere. It will also be interesting in future studies to investigate DA dynamics in further striatal subregions, i.e. assessing how the DA signal might vary across the DS from the dorsolateral to the dorsomedial part.

Upon blockage of DA reuptake by exposure to cocaine, both regions exhibited massive elevation in the DA signal on a slow time scale. For VS, this was accompanied by an almost complete loss of correlation to behavioral syllables, while the correlation between DS DA and syllables increased. This differential response was reflected in a remarkable decoupling of the dorsoventral striatal coordination as shown in the cross-correlation analysis. Combined, this might speak in favor of the previously proposed gate-like role of DA in VS (57), where the moment-to-moment DA levels constitute a signal to exert distinct behaviors under normal circumstances. When DA is artificially elevated by cocaine, it seems therefore that DA signaling in DS takes control. However, in addition to increases in basal DA, the rapid DS fluctuations no longer carried salient information based on an unchanged variance of the derivative across intensities, yet these apparently random changes in DS DA still correlated with syllables.

Curiously, this was accompanied by a more disordered sequencing of syllables as apparent from the increase in transition entropy, despite the general upregulation of the most frequent syllables. We speculate that the co-occurrence of these two phenomena represent a direct link between DA signaling in DS and the stringing together or structuring of actions on a moment-to-moment basis; that is, DS DA might modulate overall behavioral selection on a moment-to-moment basis but does not select specific actions. Taken together, the cocaine data both support a key role of DAT in governing discrete DA dynamics in DS and VS, and emphasize distinct functional roles of DA in DS and VS, that are directly reflected in differences in how information is encoded by the two regions. We should note, however, that cocaine also inhibits uptake of serotonin and norepinephrine (45), which could interfere with the results obtained.

In summary, we exploit the DA sensor dLight1.3b to reveal highly differential but cooperative DA dynamics in the DS and VS, that can be directly linked to discrete behavioral actions on time scales from hundreds of milliseconds to minutes. We find evidence of diverging modes of informational encoding that implores the need to consider regional differences when studying DA signaling. Moreover, we demonstrate the strength of a machine learning-based data analysis pipeline, which can be readily applied to studies across brain regions and neurotransmitters with genetically encoded biosensors. Indeed, this might represent a critical framework for future efforts aimed at gaining better insights into the complex pathobiology of neurological and psychiatric diseases involving dopamine and other important neurotransmitters.

## Materials and Methods

Detailed materials and methods with description of mice, DNA and AAV constructs, stereotaxic injections and implants, behavioral testing, immunohistochemistry, fiber photometry recordings and analysis, drugs, wavelet transform, mouse tracking and syllable analysis, information analysis and statistics are included in SI Appendix.

### Mice

All procedures were carried out in accordance with the institutional regulations and guidelines of University of Copenhagen and the Danish Animal Experimentation Inspectorate (License number: 2017-15-0201-01160). Male Th-Cre mice (58) were bred in house with female WT C57Bl6N mice supplied from Charles River as breeders.

### Stereotaxic injections and implants

Mice were anesthetized using isoflurane and placed in a Neurostar Stereotaxic Robot frame before injection of AAV constructs. For implant sites (DS and VS) used for subsequent fiber photometry, 300 nL AAV9-hSyn-dLight1.3b-WPREpA or AAV9-hSyn-EGFP-WPREpA at a titer of 3.0*10^12^ viral genomes/ml was injected at the target coordinate of the implant and 100 nL was injected 200 µm both below and above the target coordinate (**VS**: AP: 1.54mm, ML: -0.80mm, DV: -4.30mm; **DS**: AP: 1.18mm, ML. 1.70mm, DV: -3.00mm). For DREADD injections in the VTA, 500 nL AAV8-hSyn-hM4Di-mCherry-WPREpA was injected bilaterally at coordinates AP: -3.20mm, ML: +/-0.50mm, DV: -4.50mm and for SNc 300 nl was injected bilaterally at AP: -3.16mm, ML: -1.60mm, DV: -4.2mm and 200 nl was injected bilaterally at AP: -3.00mm, ML: +/-1.20mm, DV: -4.5mm. After injections, a 200 µm ø 0.37NA 1.25mm metal ferrule optic cannula (Doric Lenses) was slowly inserted at the target coordinate. Mice were tested at least 3 weeks after surgery.

### Behavioral Testing

Mice were tested and recorded in custom-made white arenas from the local technical workshop. For tests of responses to a novel environment and female soiled bedding, an arena measuring 4×30×15 cm with a 15 cm divider in the middle of one end and removable doors. For testing of self-paced exploratory activity before and after cocaine, we used an arena measuring 50×50×40 cm that the mice previously had been exposed to 2-3 times. Mice were recorded from above using an ELP KL36IR 1080P Full HD Webcam at 30 frames pr. second. For each test, hM4Di-injected mice were tested in a crossover design with CNO and vehicle administration. Half of the mice received CNO on the first test day and vehicle on the second while the other half received vehicle on test day one and CNO on the second test day. Data was pooled for both test days.

### Fiber photometry recordings

Fiber photometric recordings of dLight1.3b fluorescence was measured using primarily the Neurophotometrics FP3001. Some control experiments were performed on the Neurophotometrics FP3002 as the setup was updated during the course of the experiments. For all experiments, except the ones using female bedding, two mice were recorded simultaneously in separate arenas using a 3 m long multi branching fiber optic patch cord (Doric Lenses, 200 um, NA 0.37) attached to the implanted optic cannulas using bronze mating sleeves (Thorlabs ADAL 4-5). Fiber photometry recordings were performed using the open-source software Bonsai (59). Light power of the 470 nm channel was ∼25 µW measured at the patch cord tip at constant illumination (slight differences between different fibers of the patch cord). The power of the isosbestic 415 nm channel was ∼18 µW measured at the patch cord tip at constant illumination.

### Fiber photometry analysis

Raw intensity measurements were pre-processed by subtracting a technical baseline from recordings with light-impermeable caps followed by the isosbestic 415 nm reference to account for artifacts. For cocaine trials, a linear regression to correct for photobleaching was fitted to data from before vehicle injection and the last ten minutes of the trial and applied across the series. The resulting signal was converted to ΔF/F0 by dividing with the reference signal. For comparison across mice a running z-score (zF) was computed using the mean and standard deviation from the 5 minutes preceding injection (SI Appendix, Fig. S1C).

### Mouse tracking and syllable analysis

Using top-down recorded videos, mice were tracked using DeepLabCut v2.2rc3 (43). Coordinates for snout, shoulders, hips and base-of-tail were extracted and used to segregate movements into individual syllables using B-SOiD v2.0 (44). 67 clusters were identified, several empty, and due to inter-mouse variation and infrequent use, only 20 syllables were analyzed in the study.

### Statistical analysis

Choice of statistical analysis is presented in the legends associated with each figure, and multiple comparisons is corrected for using either Tukey HSD procedure for post-hoc ANOVA testing, Bonferroni-Holm when t-test are performed, or Benjamini-Hochberg false discovery rate for the information analysis due to the non-negative correlation between tests. All *n*-values are individual mice.

### Data and code availability

Data and associated codes are available at https://github.com/GetherLab/In-vivo-dopamine-dynamics. Further details of the analysis are available from the corresponding author upon request.

## ACKNOWLEDGMENTS

We thank Anette Dencker Kaas for excellent technical assistance. The work was supported by the Lundbeck Foundation grants R266-2017-4331 (UG), R276-2018-792 (UG), R230-2016-3154 (M.D.L.), R181-2014-3090 (FH), R303-2018-3540 (F.H.) and R231-2016-2481-5 (ATS), Independent Research Fund Denmark – Medical Sciences (U.G. 7016-00325B).

## Supporting Information for

### Supplementary Materials and Methods

#### Mice

All procedures were carried out in accordance with the institutional regulations and guidelines of University of Copenhagen and the Danish Animal Experimentation Inspectorate (License number: 2017-15-0201-01160). Male Th-Cre mice (1) were bred in house with female WT C57Bl6N mice supplied from Charles River as breeders. Mice were kept at a normal day cycle with light on 6-18 and were tested during the light phase. Mice had access to standard rodent chow and water ad libitum. Mice were genotyped using an appropriate PCR procedure on ear-samples. The mice underwent surgery when they were ∼12 weeks old.

#### DNA and AAV constructs

The plasmid pAAV-hSyn-dlight1.3b-WPREpA was generated by digesting pAAV-CAG-dLight1.3b (Addgene Plasmid #125560) with BamHI and HindIII restriction enzymes and cloning the resulting fragment into a pAAV-hSyn-MultipleCloningSite-WPREpA plasmid. Successful cloning was confirmed by sequencing using Mix2Seq at Eurofins Genomics. pAAVhSyn-dlight1.3b-WPREpA and pAAV-hSyn-EGFP-WPREpA (Addgene plasmid #50465) were both packaged into AAV using a previously described method (2, 3). Briefly, HEK293FT cells were triple transfected with the dLight or EGFP plasmid, rep2/cap9 plasmid and pAdDeltaF6 helper plasmid using a FuGENE 6 (Promega) transfection protocol. After 6 hours, cell medium was changed to low serum medium, and cells were harvested 3 days later. Cells were subjected to 3 freeze thaw cycles in lysis buffer and AAV particles were then isolated in an iodixanol gradient using 15-hour ultracentrifugation at 28.000 rpm in an SW28 swinging-bucket rotor, yielding a relative centrifugal field of 104,000 x g at the midpoint of the column (r_av_). The virus fraction was then harvested from the gradient using a syringe and needle and this fraction was washed and upconcentrated in Amicon filter tubes. AAVs titers were determined using a PicoGreen based assay (4) and diluted in DPBS (Sigma Aldrich #D8537) to 3.0*10^12^ viral genomes/ml before injection. AAV8-hSyn-DIO-hM4Di-WPREpA was acquired pre-packaged from Addgene (#44362-AAV8).

#### Stereotaxic injections and implants

Mice were anesthetized using isoflurane and placed in a Neurostar Stereotaxic Robot frame on top of a heating pad. A droplet of lidocaine hydrochloride was injected subdermally for local anesthesia, and eyes were covered in eye ointment (Ophta) to prevent drying. The fur on top of the scalp was removed and the surgical area was sterilized with 70 % ethanol and Povidone-Iodine 10 % using sterilized cotton applicators. An incision was made down the midline exposing bregma and lambda, and the skull was scratched using a dental drill to increase adhesion of dental cement. A droplet of 3 % hydrogen peroxide solution in sterile water was applied to clean and dry the skull for optimal dental cement adhesion. The skull was thoroughly rinsed with sterile saline to remove any residual hydrogen peroxide and dried with sterile cotton applicators. A sterilized piece of aluminum foil was placed over the eyes to prevent light induced damage from subsequent dental cement light curing. Bregma and Lambda were identified on the skull and saved in the Neurostar software. A thin layer of Optibond™ FL sealing primer (Kerr™) was applied to the skull and allowed to dry whereafter a layer of Optibond™ FL Adhesive (Kerr™) was added to the skull and light cured using a blue light gun. Injection and implant sites were identified, and holes were drilled above these using the Neurostar drill. Hereafter injection of AAV constructs began. For implant sites (DS and VS) used for subsequent fiber photometry, 300 nL AAV9-hSyn-dLight1.3b-WPREpA or AAV9-hSyn-EGFP-WPREpA at a titer of 3.0*10^12^ viral genomes/ml was injected at the target coordinate of the implant and 100 nL was injected 200 µm both below and above the target coordinate (**VS**: AP: 1.54mm, ML: -0.80mm, DV: -4.30mm; **DS**: AP: 1.18mm, ML. 1.70mm, DV: -3.00mm). For DREADD injections in the VTA, 500 nL AAV8-hSyn-hM4Di-mCherry-WPREpA was injected bilaterally at coordinates AP: -3.20mm, ML: +/- 0.50mm, DV: -4.50mm and for SNc 300 nl was injected bilaterally at AP: -3.16mm, ML: - 1.60mm, DV: -4.2mm and 200 nl was injected bilaterally at AP: -3.00mm, ML: +/-1.20mm, DV: - 4.5mm. All injections were carried out at a rate of 50-100 nl per min and the glass needle was left in place for 5 min after injection to allow diffusion into the tissue before being slowly removed. After injections, a 200 µm ø 0.37NA 1.25mm metal ferrule optic cannula (Doric Lenses) was slowly inserted at the target coordinate. The metal ferrule was attached to the skull using light curing Tetric EvoFlow (Ivoclar Vivadent) dental cement. The incision was closed using absorbable Vicryl suture (Ethicon), and mice were given 1 ml saline s.c. to aid rehydration after surgery. Mice were allowed to wake up in individual cages before returning to their home cage with their littermates. Mice were treated with an analgesia/antibiotic mixture of carprofen (Rimadyl) and enrofloxacin (Baytril) in saline on the day of surgery and the two following days. Mice were tested at least 3 weeks after surgery.

#### Behavioral Testing

Mice were tested and recorded in custom-made white arenas from the local technical workshop. For tests of responses to a novel environment and female soiled bedding, an arena measuring 45×30×15 cm with a 15 cm divider in the middle of one end and removable doors. These doors were removed manually at pre-determined timepoints to allow access to the closed compartments. Female soiled bedding was collected and combined from two separate female cages, divided into equal portions, and placed in one of the closed compartments in lids of 50 ml centrifuge tubes. For testing of self-paced exploratory activity before and after cocaine, we used an arena measuring 50×50×40 cm that the mice previously had been exposed to 2-3 times. Mice were recorded from above using an ELP KL36IR 1080P Full HD Webcam at 30 frames pr. second. Arenas were illuminated using flicker-free light bulbs (Toshiba e27 60W) to prevent banding effects in the videos. For each test, hM4Di-injected mice were tested in a crossover design with CNO and vehicle administration. Half of the mice received CNO on the first test day and vehicle on the second while the other half received vehicle on test day one and CNO on the second test day. Data was pooled for both test days.

#### Immunohistochemistry

After finalizing experiments, mice were anaesthetized using isoflurane and transcardially perfused with cold PBS (phosphate-buffered saline) and subsequently 4% PFA (paraformaldehyde) in PBS Brains were removed and dropped in 4% PFA overnight. Brains were then transferred to 30 % sucrose in PBS and left for two days, whereafter the brains were frozen on dry ice and stored at -80° C. Brains were cut in 40 µm thick slices in a cryostat. Slices of interest were stained for presence of the dLight1.3b using an anti-GFP antibody (Abcam - ab13970, 1:1000). No enhancement was needed to visualize hM4Di-mCherry expression. After washing in PBS 3 times 5 minutes slices were pre-incubated for 30 min in a PBS solution containing 5% goat serum, 1% BSA and 0.3% Triton X-100 at room temperature. Slices were then transferred to the same solution but containing the primary antibody and left to incubate at 4° C overnight. Slices were then washed 3 times 5 minutes in a PBS solution containing 0.25 % BSA and 0.1 % Triton X- 100 and hereafter incubated for 1 hour in an identical solution containing secondary antibody (Alexa 488, goat-α-chicken, Ab150173, 1:400). Hereafter, slices were mounted on glass slides, allowed to air-dry and covered with a coverslip using prolong gold antifade mounting medium (Thermo Fisher). Slices were inspected and probe locations were verified using an epifluorescent microscope (Zeiss Axiovert 100 or Leica DM IL LED was used) and representative images were acquired using a spinning disk confocal microscope at 10- and 20-times magnification (FEI CorrSight, Core Facility for Integrated Microscopy -University of Copenhagen). Images were stitched using the ImageJ plugin MIST and the linear stitching method {Chalfoun, 2017 #56}.

#### Fiber photometry recordings

Fiber photometric recordings of dLight1.3b fluorescence was measured using primarily the Neurophotometrics FP3001. Additionally, some control experiments were performed on a Neurophotometrics FP3002 as the setup was updated during the course of the experiments. For all experiments, except the ones using female bedding, two mice were recorded simultaneously in separate arenas using a 3 m long multi branching fiber optic patch cord (Doric Lenses, 200 um, NA 0.37) attached to the implanted optic cannulas using bronze mating sleeves (Thorlabs ADAL 4-5). Fiber photometry recordings were performed using the open-source software Bonsai (6). Light power of the 470 nm channel was ∼25 µW measured at the patch cord tip at constant illumination (slight differences between different fibers of the patch cord). The power of the isosbestic 415 nm channel was ∼18 µW measured at the patch cord tip at constant illumination. The power of the 415 nm channel was set slightly lower than the 470 nm channel as this was found to alleviate faster bleaching of the 415 nm channel compared to the 470 nm channel, while also yielding similar absolute values for the 415 nm and 470 nm channels during recording. Light power output was measured using a light power meter (Thorlabs PM100D + S130C 400-1100 nm sensor). The same patch cord fiber was used for the same region in the same animal throughout all experiments – except for a fiber-switch-test to confirm that signal was independent of which patch cord fiber that was used.

#### Drugs

Clozapine N-oxide (CNO) (Tocris, #4936) was dissolved in DMSO (Sigma Aldrich) and then diluted in sterile saline to a final concentration of 0.4 % DMSO and 0.2 mg/ml CNO and administered by intraperitoneal injection at 10 µL/g bodyweight for a final dose of 2.0 mg/kg bodyweight. For vehicle controls, a sterile saline solution containing 0.4 % DMSO was used. Cocaine hydrochloride (Sigma Aldrich, #C-5776) was dissolved in sterile saline at a concentration of 2mg/ml and frozen in aliquots until the testing day. It was administered by intraperitoneal injection at 10 µL/g bodyweight for a final dose of 20 mg/kg bodyweight.

#### Wavelet Transform

Recorded signal (20 Hz) was split with a multiresolution analysis of the maximal overlap discrete wavelet transform at level 1 (10 Hz) and level 10 (0.0195 Hz) using the *modtwt()* and *modtwtmra()* functions of MATLAB R2018b. Signal above 10 Hz was filtered as noise as per the Nyquist theorem, signal between 10 and 0.0195 Hz as the rapid domain and anything below 0.0195 Hz taken as slow oscillations.

#### Mouse tracking and syllable analysis

Using top-down recorded videos, mice were tracked using DeepLabCut v2.2rc3 (7). Coordinates for snout, shoulders, hips and base-of-tail were extracted and used to segregate movements into individual syllables using B-SOiD v2.0 (8). 67 clusters were identified, several empty, and due to inter-mouse variation and infrequent use, only 20 syllables were analyzed in the study.

#### Random forest classifier

The random forest classifier was built using the open source scikit-learn v0.24.1 (9). The model was trained on Nx402 linearized data of DS and VS signal and split 80/20 for training and test. Hyper parameters were tuned using the RandomizedSearchCV module and set to 2,500 trees, depth of 30 and minimum sample split and leaf size of 2.

#### Signal heterogeneity analysish

Heterogeneity of the fiber photometry signal were assessed by highpass filtering the slow domain, truncating the upper and lower 0.1% of the signal, segmenting the remainder into deciles and computing the variance of the first order derivative for each segment. Entropy rate was calculated from the transition matrix of each mouse using the method described in Wiltschko et al., Neuron 2015 (10).

#### Statistical analysis

Choice of statistical analysis is presented in the legends associated with each figure, and multiple comparisons is corrected for using either Tukey HSD procedure for post-hoc ANOVA testing, Bonferroni-Holm when t-test are performed, or Benjamini-Hochberg false discovery rate for the information analysis due to the non-negative correlation between tests. All *n*-values are individual mice. Statistical analyses were carried out with the open-source python packages SciPy v1.5.2 (11), NumPy v1.18.1 (12), and Seaborn v0.11.0 (13), and linear models in Statsmodels v0.12.2. Boxplots show 25^th^ and 75^th^ percentile, with whiskers indicating data within 1.5 times the interquartile range. Remaining data were plotted as outliers. No statistical methods were used to predetermine sample sizes. Mice were randomly allocated to different groups for the in vivo experiments, but data collection was not performed under blind conditions. The analyses were not performed under blinded conditions but were automated to a degree where the experimenter had no impact on outcome. Single animals were excluded from data analysis after tissue analysis had confirmed no or off-target viral expression. Sessions with severe fiber tangling were excluded from analysis on a qualitative basis.

## Supporting Figures

**Fig. S1.**
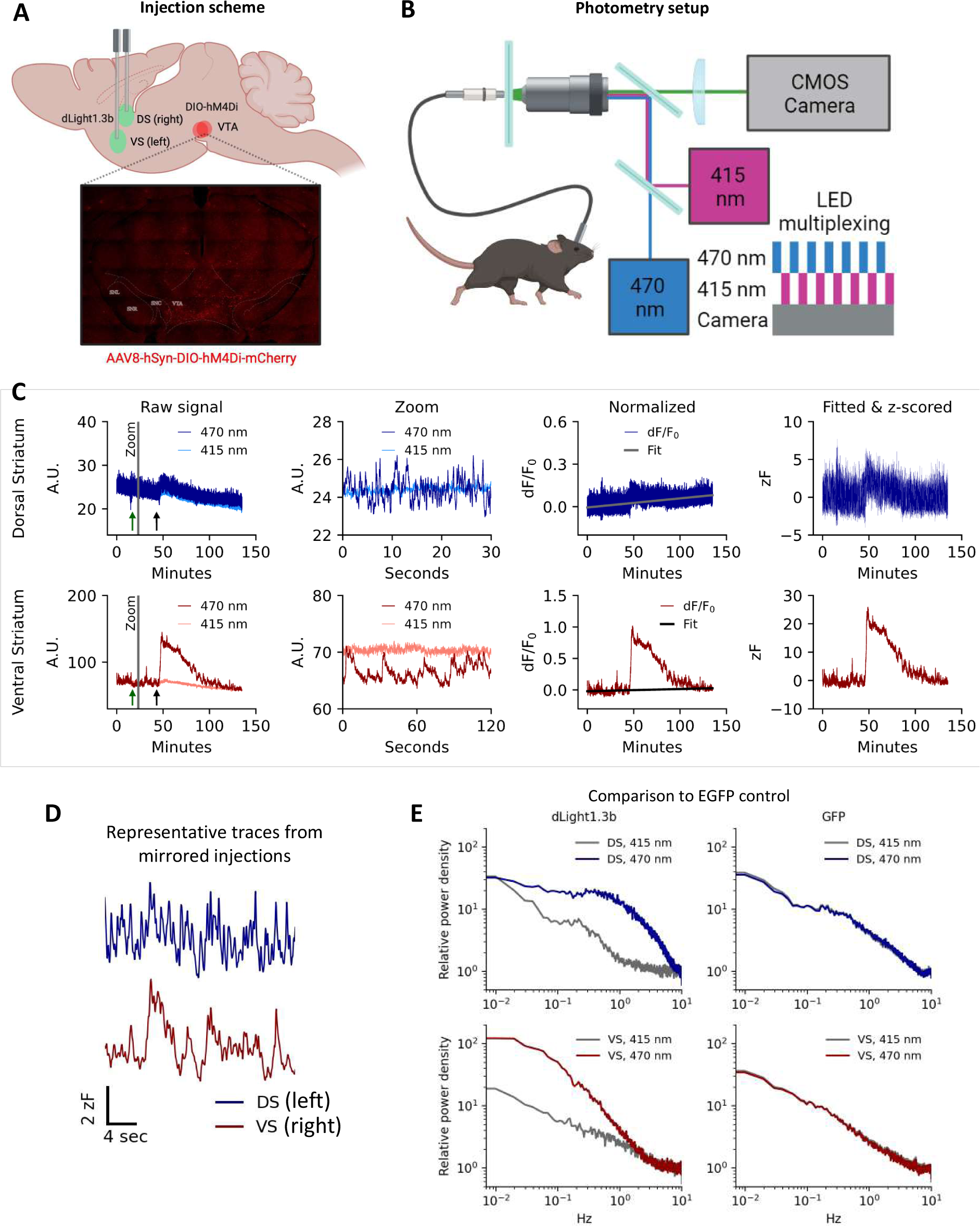
Injections and schematics. (A) Injection and implantation scheme for dLight1.3b and hSyn-DIO-hM4Di-mCherry in VTA in TH-Cre mice. Inserted image shows histological validation of mCherry expression in VTA. (B) Schematic representation of the fiber photometry system. (C) Procedure for signal pre-processing. Left panels, Raw signal from 415 and 470 nm channels for DS (top) and VS (bottom) in representative mouse. Green arrow indicates injection of vehicle, black arrow indicates injection of cocaine (20 mg/kg, i.p.). Middle left; Zoom (indicated with a gray line in left panels) on the rapid pre-injection fluctuations of the raw signal from 415 and 470 nm channels for DS (top) and VS (bottom). Middle right, The two channels are converted to dF/F_0_, and a linear fit is applied to the entire trace based on data from before vehicle injection and the last ten minutes of the trial to correct for photobleaching. Right, Signal is z-scored (zF) to the pre-injection variation to facilitate comparison across mice. A.U. = arbitrary units. (D) Representative traces of DA fluctuations in left DS (blue) and right VS (red) as assessed by dLight1.3b fluorescence (expression of dLight1.3b mirrored compared to data shown in Fig. 1). (E) Representative power spectrum for 415 nm and 470 nm channels in fiber photometry recordings of WT mice injected with AAV encoding either dLight1.3b or GFP. While a clear signal was seen for dLight1.3b injected mice in both DS and VS, we observed only noise when mice were injected with EGFP in both regions (data are representative of three different GFP expressing mice).

**Fig. S2.**
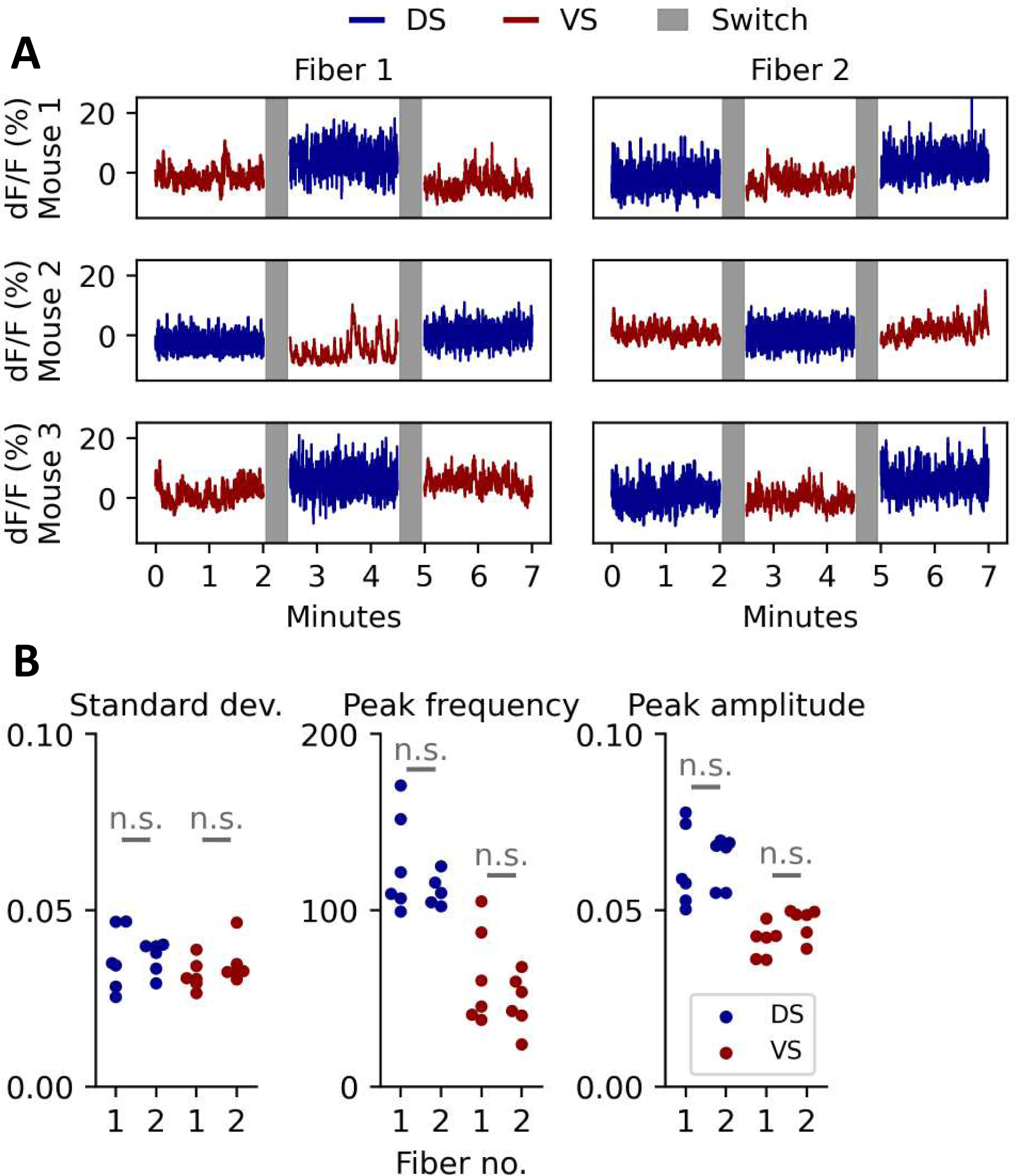
Switching the recording patch cord fibers mid-session robustly produces the same result for both regions. (A) Representative traces of dLight1.3b fluorescence in DS and VS from three different mice where the recording patch cord fibers are switched after ∼2 min and then switched back ∼2 min later. (B) Quantifying standard deviation, peak frequency and peak amplitude of the signal recorded in A shows no statistical difference between the two patch cord fibers (standard deviation: DS, p = 1.0; VS, p = 0.93; peak frequency: DS, p = 1.0; VS, p = 0.84; peak amplitude: DS, p = 1.0; VS, p = 0.18; paired, student’s t-test, FWER correction by region with Bonferroni-Holm).

**Fig. S3.**
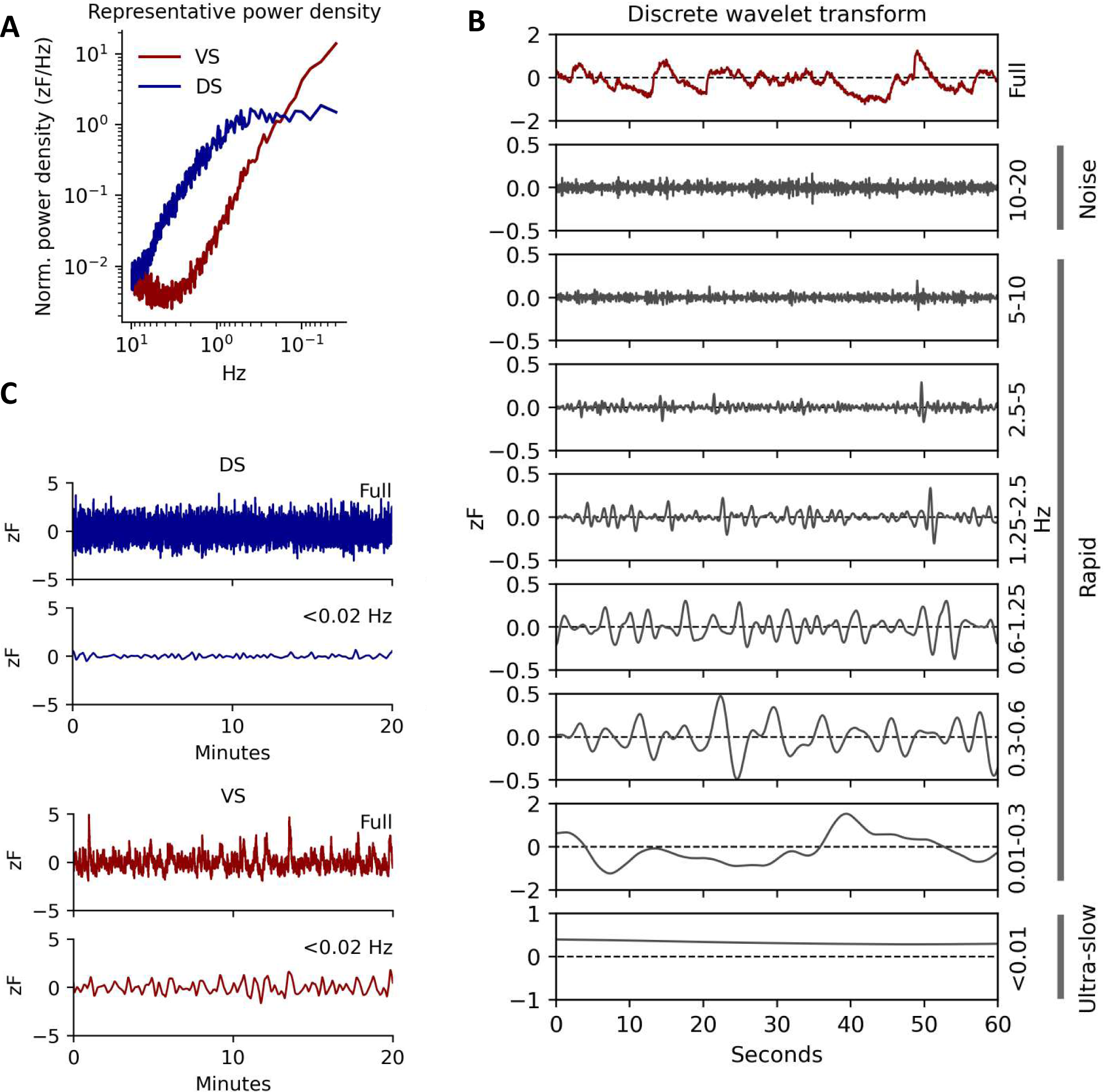
Wavelet transform and full cross-correlation. (A) A representative power density of the dynamics of the dLight1.3b fluorescent signal in DS (blue) and VS (red). (B) Representative example of the discrete wavelet transform split. Full signal from VS plotted in the first panel with isolated frequency bands in descending order below. By the Nyquist criterion 10 to 20 Hz is filtered as noise. Anything between 10 and 0.02 Hz is considered the rapid domain, with the remaining signal summed on the final panel and categorized as ultra-slow oscillations. (C) Representative 20-minute traces of full signal and the slow domain (<0.02 Hz) for DS (Top, dark blue) and VS (Bottom, dark red).

**Fig. S4.**
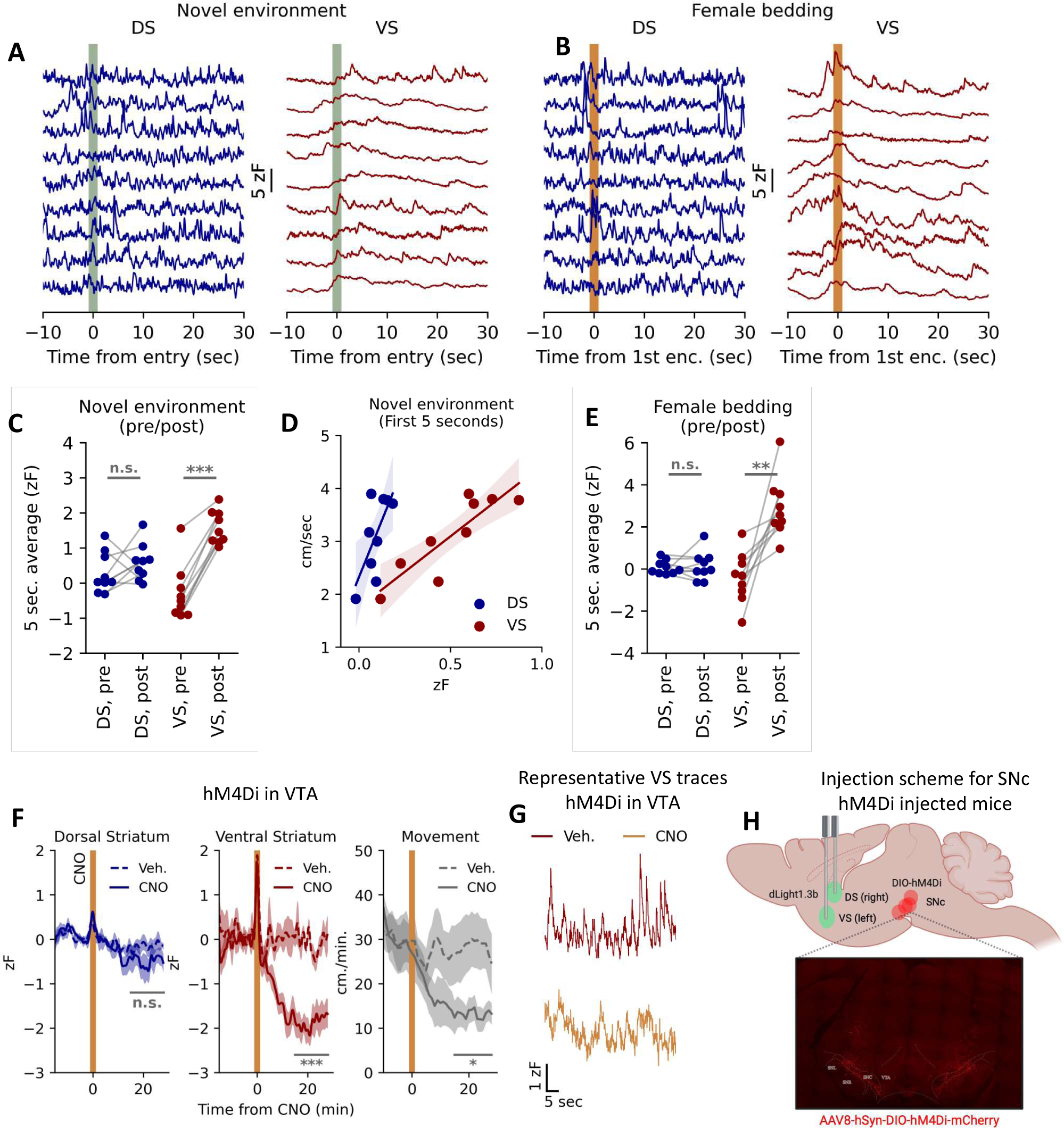
Extended data on behavioral paradigm. (A) All DS and VS fluorescence traces for the individual mice upon entry to the novel environment. Entry indicated by a dark-green band and time-locked to t = 0. (B) All DS and VS traces for the individual mice at encounter with female bedding. Time of encounter indicated by a dark-green band and time-locked to t = 0. (C) Average zF five seconds before and after entry to novel environment. Two-sided t-test, DS: p = 0.31, VS: ***p = 5.8E-06, n = 9, corrected for multiple comparisons with Bonferroni and Holm (BH) procedure. (D) zF vs. mean speed first five seconds after novel environment entry. Linear regression, slope = 0, DS: *p = 0.038, VS: **p = 0.002, n = 9, corrected for multiple comparisons with BH procedure. (E) Average zF -10 to -5 seconds before and 0 to 5 seconds after novel environment. Two-sided t-test, DS: p = 0.78, VS: **p = 0.004, n = 9, corrected for multiple comparisons with BH procedure. (F) Fluorescent DA trace in DS (left) and VS (middle) and the parallel locomotion (right) after i.p. injection of either vehicle or 2 mg/kg CNO in TH-Cre mice expressing dLight1.3b in the right DS and left VS, and AAV-hSyn-DIO-hM4Di-mCherry in VTA; vehicle (dashed line) and CNO (solid line). All traces are time-locked to injection (t = 0, orange band). As assessed by area under the curve, the difference between vehicle and CNO was not significant for DS (n.s., p = 0.21) but significant for VS (***p = 8.3E-5). Movement was significantly decreased (*p = 0.03); unpaired, two-tailed student’s t-test. Shaded area indicates S.E.M., n = 9 for both groups. (G) Representative trace for vehicle (dark red) and CNO (orange). (H) Despite lack of effect of CNO administration, we observed clear DREADD expression in SNc. Images visualize hM4Di-mCherry expression in TH-Cre mice stereotactically injected with AAV-hSyn-DIO-hM4Di-mCherry bilaterally in SNc, two viral injections per hemisphere were used to target SNc.

**Fig. S5.**
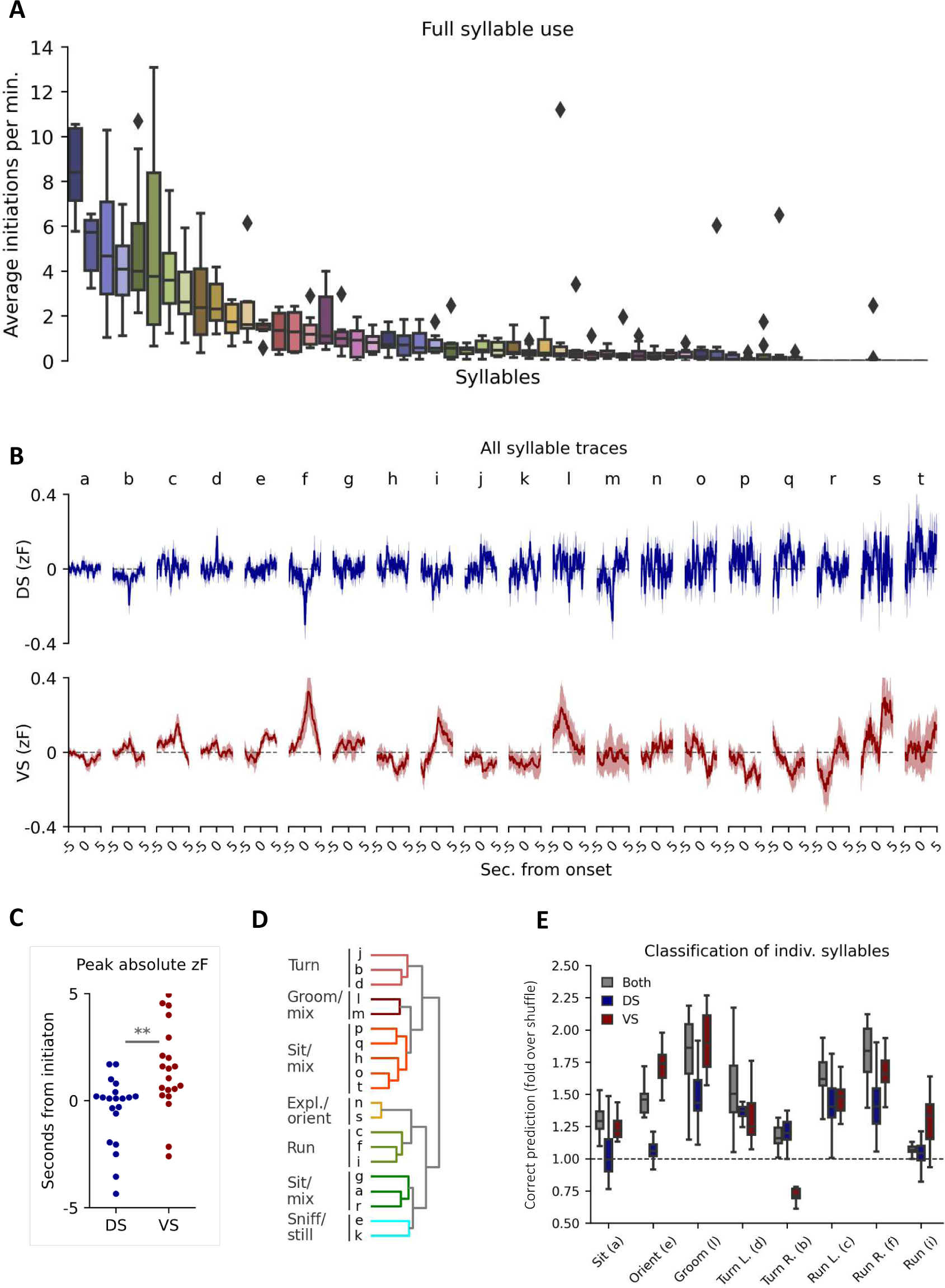
Extended syllable overview. (A) Box plots of average number of initiations per minute across all identified syllables (<50) (n = 9 mice). The top 20 syllables were chosen for further analysis. (B) Average fluorescent DA trace at onset for DS (blue) and VS (red) for top 20 syllables. Shaded areas indicate S.E.M., n = 9. (C) Comparison of maximal absolute value of the signal (zF) surrounding syllable initiation for DS and VS (traces in Fig. S5B), two-sided t-test, **p = 0.0058. (D) Human annotations for seven groups of syllables identified by hierarchical clustering of Spearman correlation coefficients. (E) Box plots of correct prediction rate (fold over shuffle) for eight most unambiguous syllables, shown for combined traces, or only DS or VS traces.

**Fig. S6.**
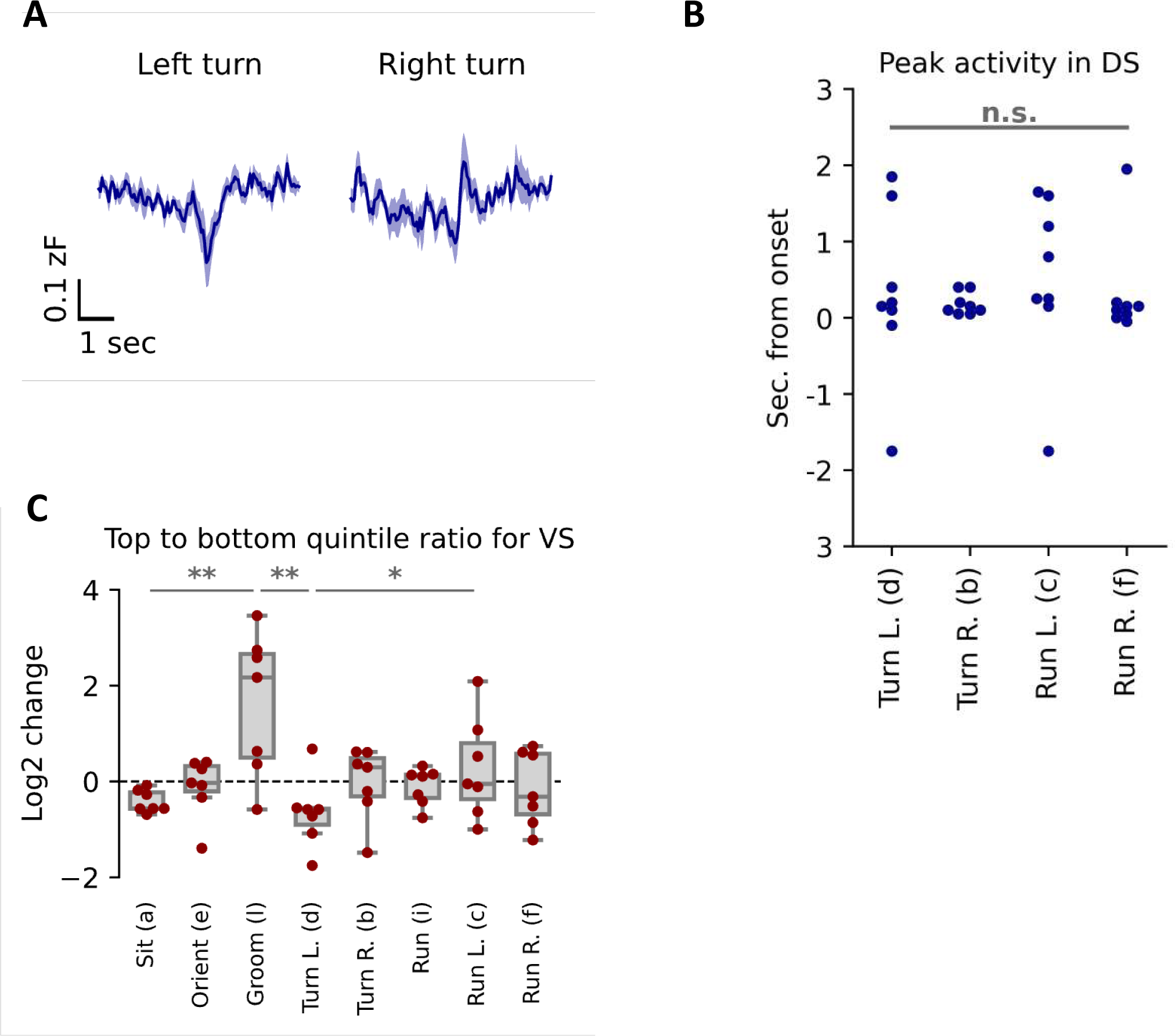
(A) Expression of dLight1.3b in the opposite hemisphere (left DS) showed a mirror of the signal seen in the right DS (Fig. 3E). Average fluorescent DA trace around onset in left DS for left and right turn syllables (n=9). Shaded area indicates S.E.M. Due to a new video recording setup, tracking and unsupervised clustering was redone as described in Methods. (B) Timing of peak DS activity during turns. For neither stationary nor running turns max [DA] timing differs statistically from syllable onset (turn L., p = 1.0; turn R., p = 0.62; run L., p = 1.0; run R., p = 1.0. student’s t-test, H0 = 0, n = 9, FWER correction by domain with Bonferroni-Holm) (n=9). (C) Box plot of difference in frequency for upper and lower quintile of slow VS levels, as conceptualized on Fig. 3F (one-way ANOVA for all 8 syllables in Fig. 3E, F = 3.5, **p = 0.004, n = 9. Post-hoc Tukey HSD: a = l, **p = 0.01, l = d, **p = 0.008, d = c, *p = 0.03)

**Fig. S7.**
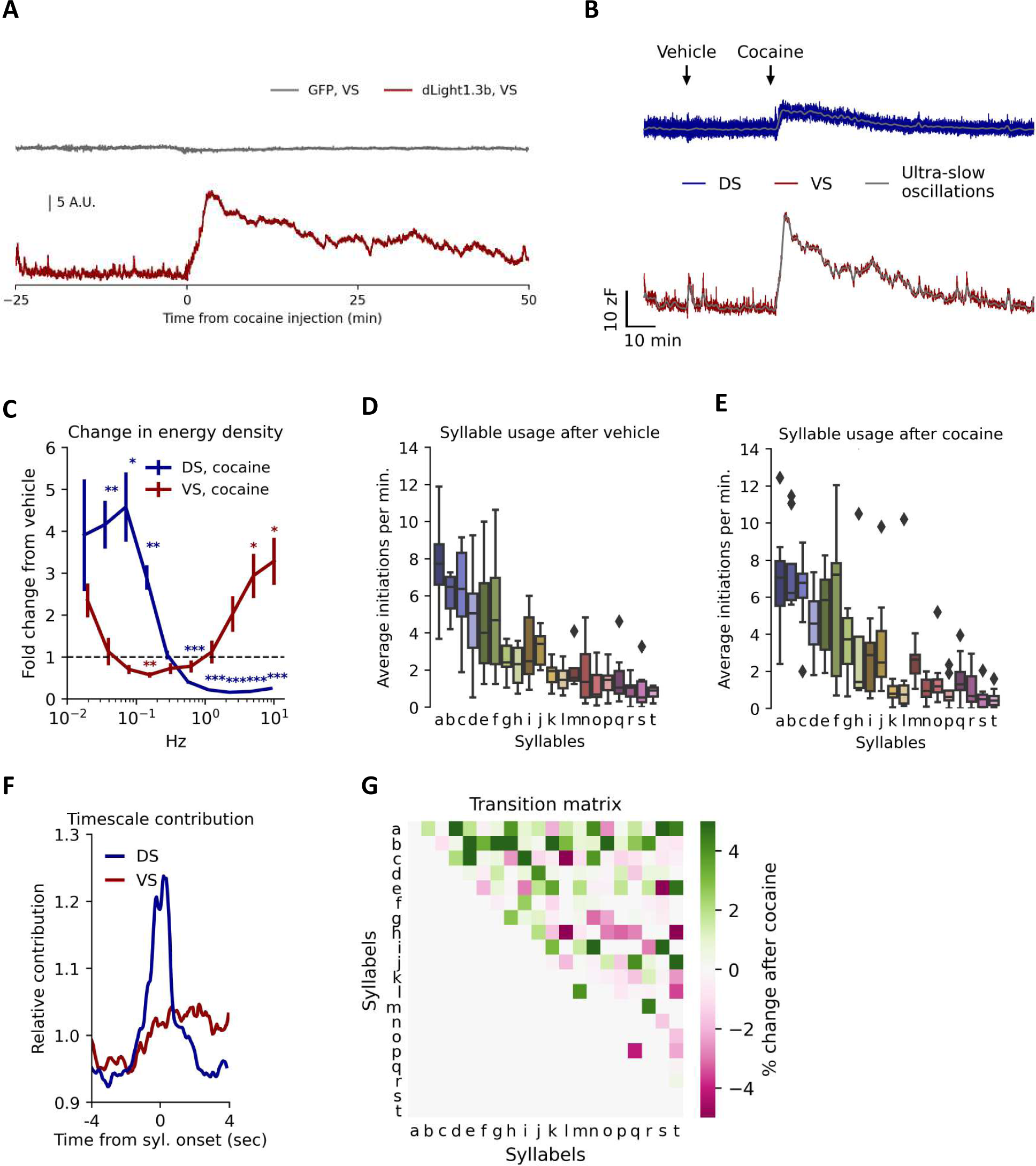
Cocaine trace and syllable use. (A) Representative fluorescent traces in VS showing response to vehicle or cocaine for a representative mouse injected with either GFP or dLight1.3b (representative of three different GFP expressing mice). Cocaine did not elicit any change in the signal from mice expressing GFP but caused a substantial increase in the dLight1.3b signal. (B) Representative fluorescent DA traces in DS (blue) and VS (red) after vehicle and cocaine administration. Wavelet-isolated ultra-slow oscillations (<0.01 Hz) depicted in grey. (C) Change in spectral energy density of the fluorescent DA signal from saline to cocaine (10 min after administration) quantified for both DS (dark blue) and VS (dark red). Error bars indicate S.E.M., n = 9, student’s t-test, H_0_ = 1, FWER correction for all ten frequency bands with Bonferroni-Holm method. (D) Box plot of syllable usage for 20 most frequent syllables after vehicle administration, n = 9. (E) Box plot of syllable usage for 20 most frequent syllables after cocaine administration, n = 9. (F) Relative contribution to classifier prediction during cocaine by region (DS, blue. VS, red) around onset of syllables. (G) Syllable transition matrix showing absolute percentage changes between vehicle and cocaine treatment.

## Notes

### Summary of Updates

Small error proofs

